# Genomic basis of European ash tree resistance to ash dieback fungus

**DOI:** 10.1101/626234

**Authors:** Jonathan J. Stocks, Carey L. Metheringham, William Plumb, Steve J. Lee, Laura J. Kelly, Richard A. Nichols, Richard J. A. Buggs

## Abstract

Populations of European ash trees (*Fraxinus excelsior*) are being devastated by the invasive alien fungus *Hymenoscyphus fraxineus*, which causes ash dieback (ADB). We sequenced whole genomic DNA from 1250 ash trees in 31 DNA pools, each pool containing trees with the same ADB damage status in a screening trial and from the same seed-source zone. A genome-wide association study (GWAS) identified 3,149 single nucleotide polymorphisms (SNPs) associated with low versus high ADB damage. Sixty-one of the 203 most significant SNPs were in, or close to, genes with putative homologs already known to be involved in pathogen responses in other plant species. We also used the pooled sequence data to train a genomic prediction (GP) model, cross-validated using individual whole genome sequence data generated for 75 healthy and 75 damaged trees from a single seed source. Using the top 30% of our genomic estimated breeding values from 200 SNPs, we could predict tree health with over 90% accuracy. We infer that ash dieback resistance in *F. excelsior* is a polygenic trait that should respond well to both natural selection and breeding, which could be accelerated using GP.

## Introduction

*Fraxinus excelsior* (European ash), is a broad-leaved tree species widespread in Europe, with over 900 dependent species^1,2^, and with high genetic diversity^3^. Its populations are being severely reduced by the invasive alien fungus *Hymenoscyphus fraxineus*, which causes ash dieback^4^. Several previous studies have shown that there is a low frequency of heritable resistance to ADB in European ash populations^5^. Estimates of breeding values of mother trees based on observed ADB damage in their progeny have an approximately normal distribution, hinting that resistance is a polygenic trait^6^ that would respond well to selection. However, an associative transcriptomics study on 182 Danish ash trees found expression levels of 20 genes associated with ADB damage scores but no genomic SNPs^3^. In model organisms, crops and farm animals, analysis of genomic information has been widely used to discover candidate genes involved in phenotypic traits, or to identify individuals with desirable breeding values^7–13^. The identification of candidate loci typically makes use of genome-wide association studies (GWAS) whereas genomic prediction (GP) methods can be used to select individuals with high breeding values. These methods have seldom been applied to keystone species in natural ecosystems due to the typically high genetic variability of such species and the high cost of genome-wide genotyping. Previous studies have demonstrated that estimation of allele frequencies by sequencing of pooled DNA samples (pool-seq) can reduce the cost of a GWAS^14^, but thus far such data have not been applied to the training of GP models. Here, we applied pool-seq GWAS and pool-seq trained GP models to European ash populations, finding a large number of SNPs associated with ADB damage that allow us to make accurate estimates of breeding values.

## Results

### Genome-wide association study

For 1250 ash trees we generated average genome coverage of 2.2x per tree, within DNA pools of 30-58 trees (Table S1). Each pool contained DNA from trees from one of thirteen seed source zones, and from trees that were either healthy or highly damaged by ADB in a mass screening trial^15^ (Figure S1, Tables S2). On average 98.3% of reads per pool mapped to the ash reference genome assembly^3^. After filtering read alignments for quality, coverage, indels and repeats, we calculated allele frequencies at 9,347,243 SNP loci. A correspondence analysis (CA), on the major allele frequencies for all 31 pools showed a distribution reflecting the geographic origin of the seed sources (Figure 1), in which axis 1 (summarising 10% of variation) reflected latitude and axis 2 (summarising 9% of variation) reflected longitude. Allele frequency measures were highly correlated in technical and biological replicates (Figure S2). In a GWAS of allele frequencies in healthy versus ADB-damaged pools, we found 3,149 significant SNPs using a Cochran-Mantel-Haenszel (CMH) test and a local FDR cut-off at 1× e^−4^ (Table S3, Figure S3). Imposing a more stringent cut-off of 1 × e^−13^, we found 203 SNP loci significantly associated with ash dieback damage scores (Figure 2).

**Figure 1.**
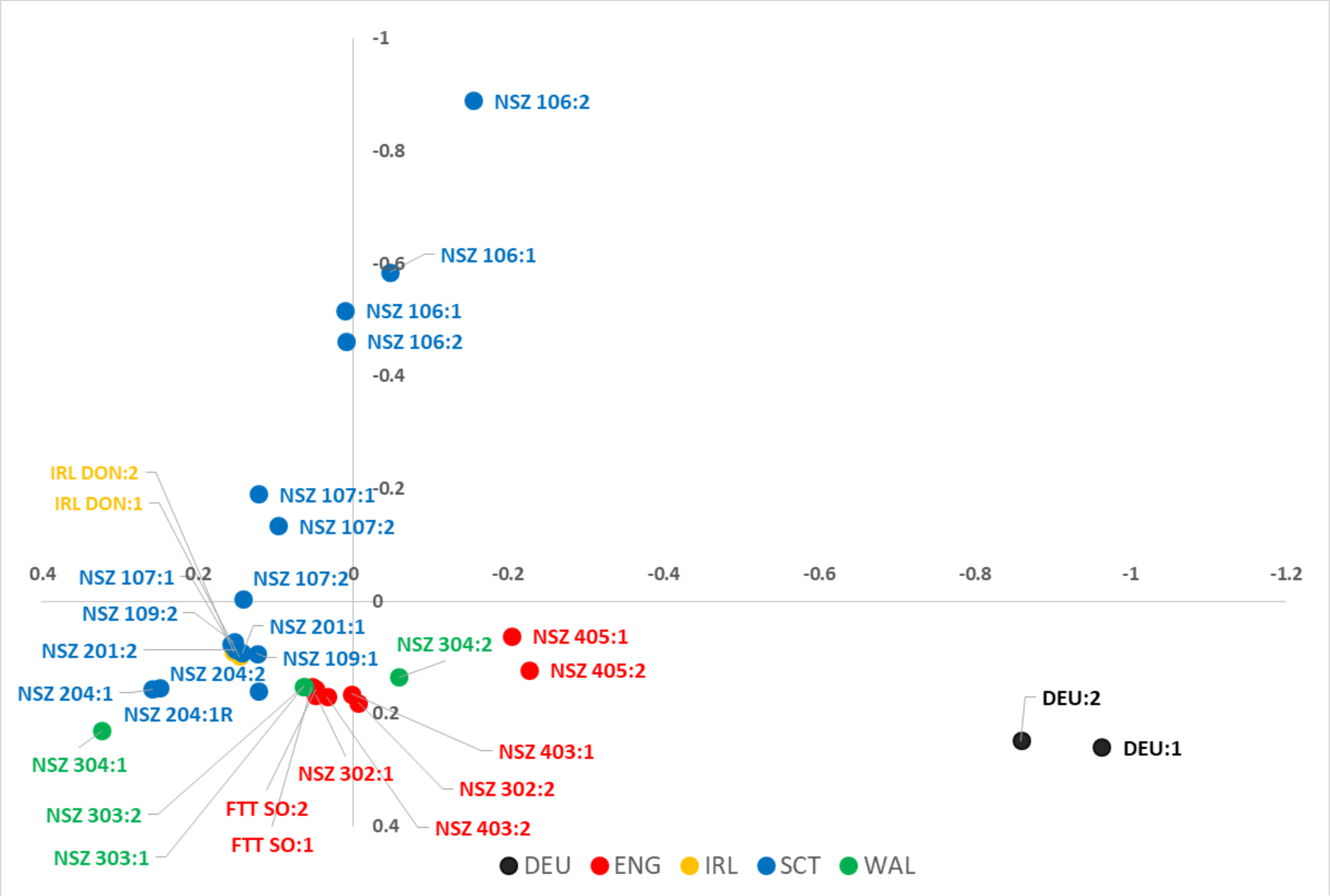
Correspondence Analysis (CA) using major allele frequency for all 31 seed source populations (including replicate). Numbers after seed source code correspond to health status (1 - healthy or 2 - infected by ADB). The vertical axis represents Principal Coordinate 1, which accounts for 10% of the variation and the horizontal axis represents Principal Coordinate 2, which accounts for 9% of the variation.

**Figure 2.**
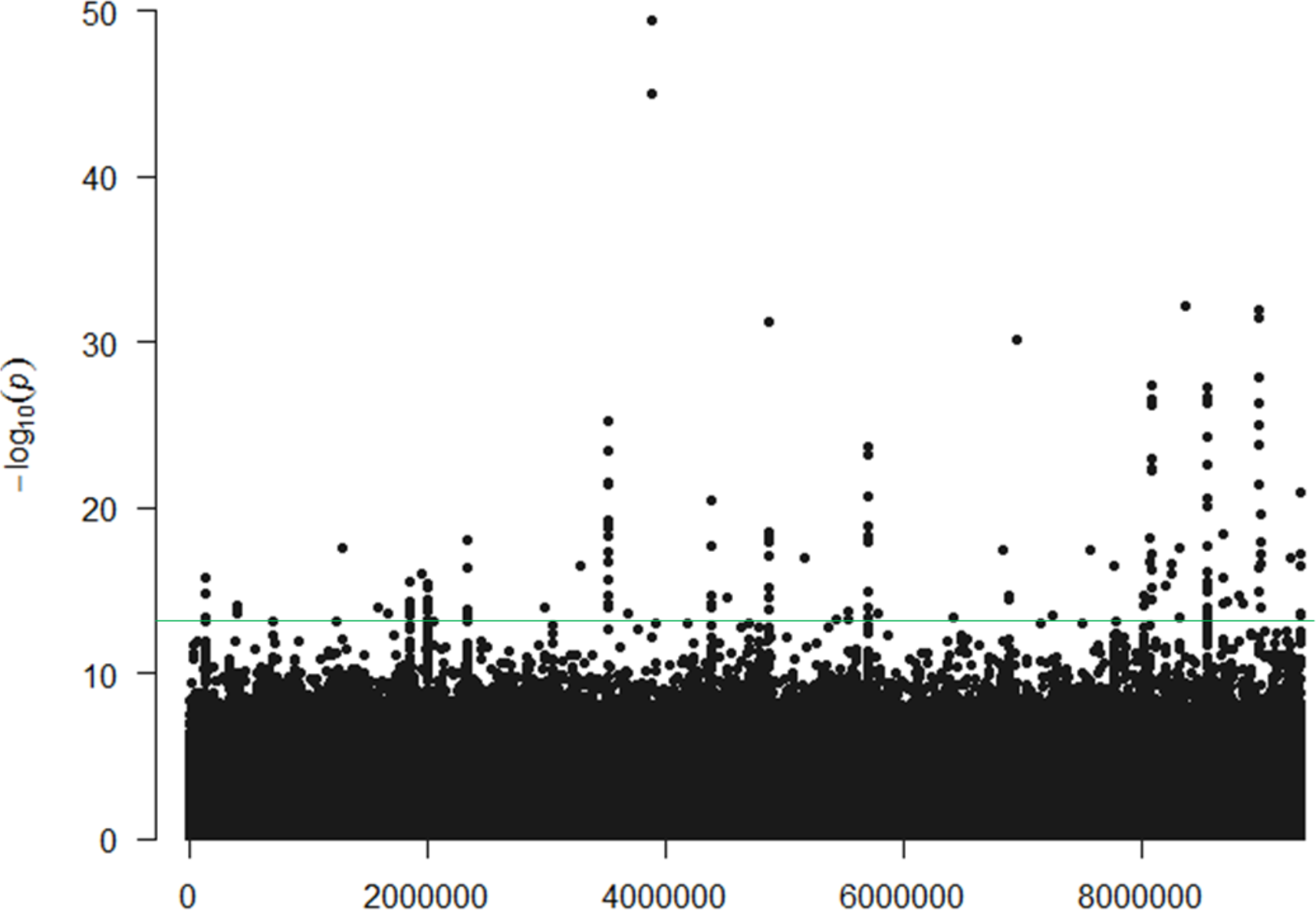
Loci associated with ash tree health status under ash dieback pressure. Genome-wide association study on whole genome sequence data from pooled DNA: Manhattan plot distribution of −log_10_(p) values for each SNP, ordered by scaffold/contig. A threshold of p = 1 × e^−13^ is shown.

Seven genes contained missense variants caused by ten of these 203 SNPs (Table 1, Figure S4, Table S5). We were able to model the proteins encoded by four of these genes (Figure 3). Similarity searches on these seven genes suggested that four of them are already known to be involved in stress or pathogen responses in other plant species. Gene FRAEX38873_v2_000003260, is putatively homologous to an *Arabidopsis* BED finger-NBS-LRR-type Resistance (R) gene (At5g63020)^16^ and is affected by a leucine/tryptophan variant close to the protein’s nucleotide binding site (Figure 3a) with the tryptophan being rarer overall, but at a higher frequency in the healthy than the damaged trees (Table S5). This R gene is located (see Figure S4) on Contig 10122 less than 5Kb from gene FRAEX38873_v2_000003270, which is putatively homologous to a Constitutive expresser of Pathogenesis-Related genes-5 (CPR5)-like protein and affected by an isoleucine/serine variant, a 5’ UTR start codon variant and 16 non-coding variants. This CPR5-like gene is likely to regulate disease responses via salicylic acid signalling^17^. Gene FRAEX38873_v2_000164520 is a putative F-box/kelch-repeat protein SKIP6 homolog, which encodes a subunit of the Skp, Cullin, F-box containing (SCF) complex, catalysing ubiquitination of proteins prior to their degradation^18^. One of our candidate SNPs encodes an arginine/glutamine substitution in this gene, with the arginine being rarer overall, but at a higher frequency in the healthy than the damaged trees. The substitution is located close to the gene’s F-box motif (Figure 3b) and is likely to affect binding within the SCF complex due to the charge difference between the two amino acids. In pine trees, F-Box-SKP6 proteins have been linked to fungal resistance^19^. Gene FRAEX38873_v2_000305440, may also be involved in ubiquitination: although the CDS hit an uncharacterised gene in olive (Table 1), the mRNA hit an E3 ubiquitin-protein ligase. This gene contains a glycine to aspartic acid substitution.

**Table 1.**
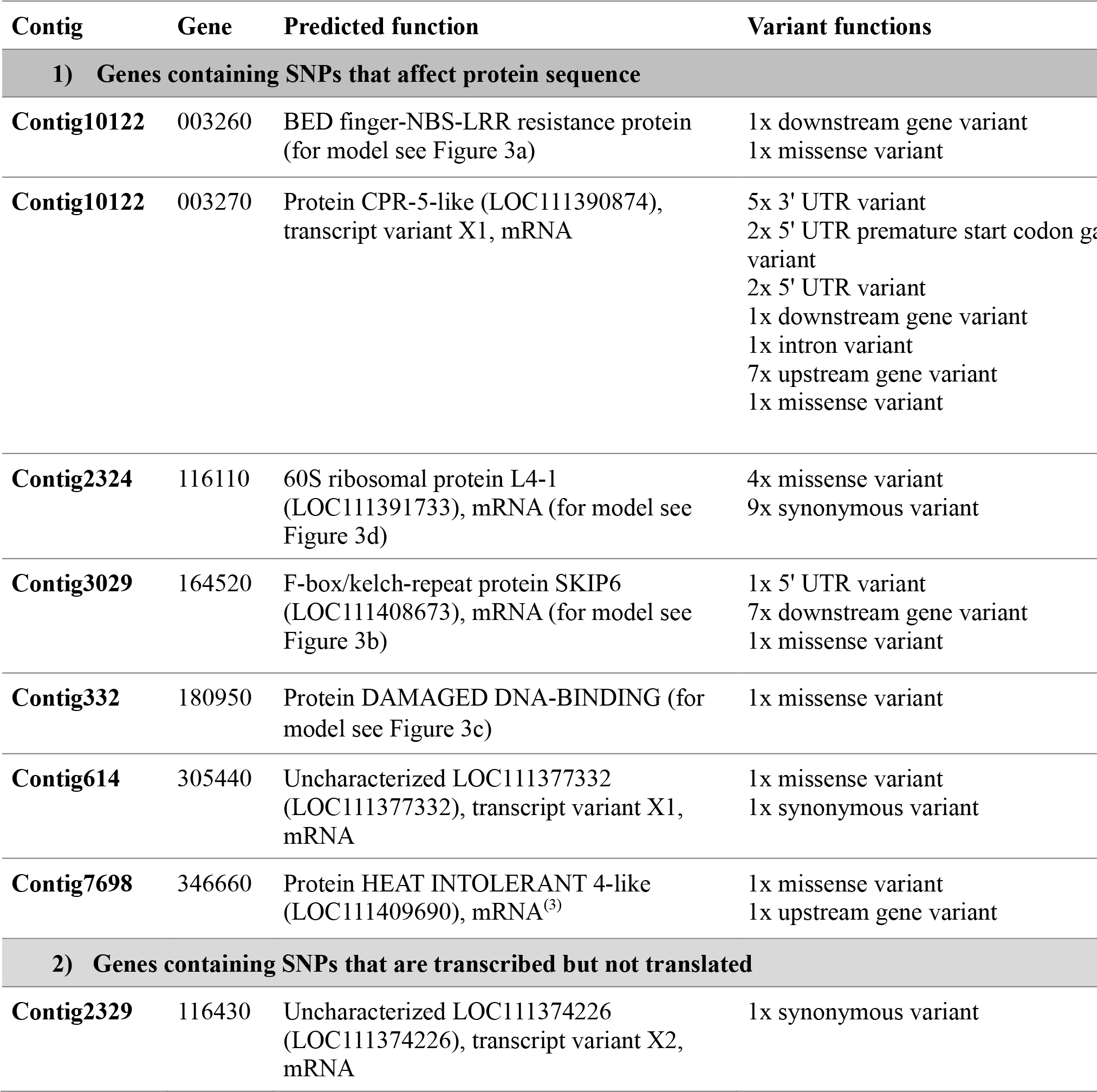
List of ash genes likely to be affected by GWAS candidate SNPs found in the top 203 hits by p-value (with −log_10_(p) > 13): (1) Genes that contain one or more significant SNP loci altering protein sequence; (2) Genes containing SNPs that are transcribed but not translated (synonymous changes, and changes in UTRs and introns); (3) Genes that are within 5Kb of significant SNP loci and the closest gene to those loci. The “Gene” column gives the final six digits for the full gene names for the annotation of the ash genome^3^, which are in the form FRAEX38873_v2_000######. Details of amino acid changes in missense variants can be found in Table S5.

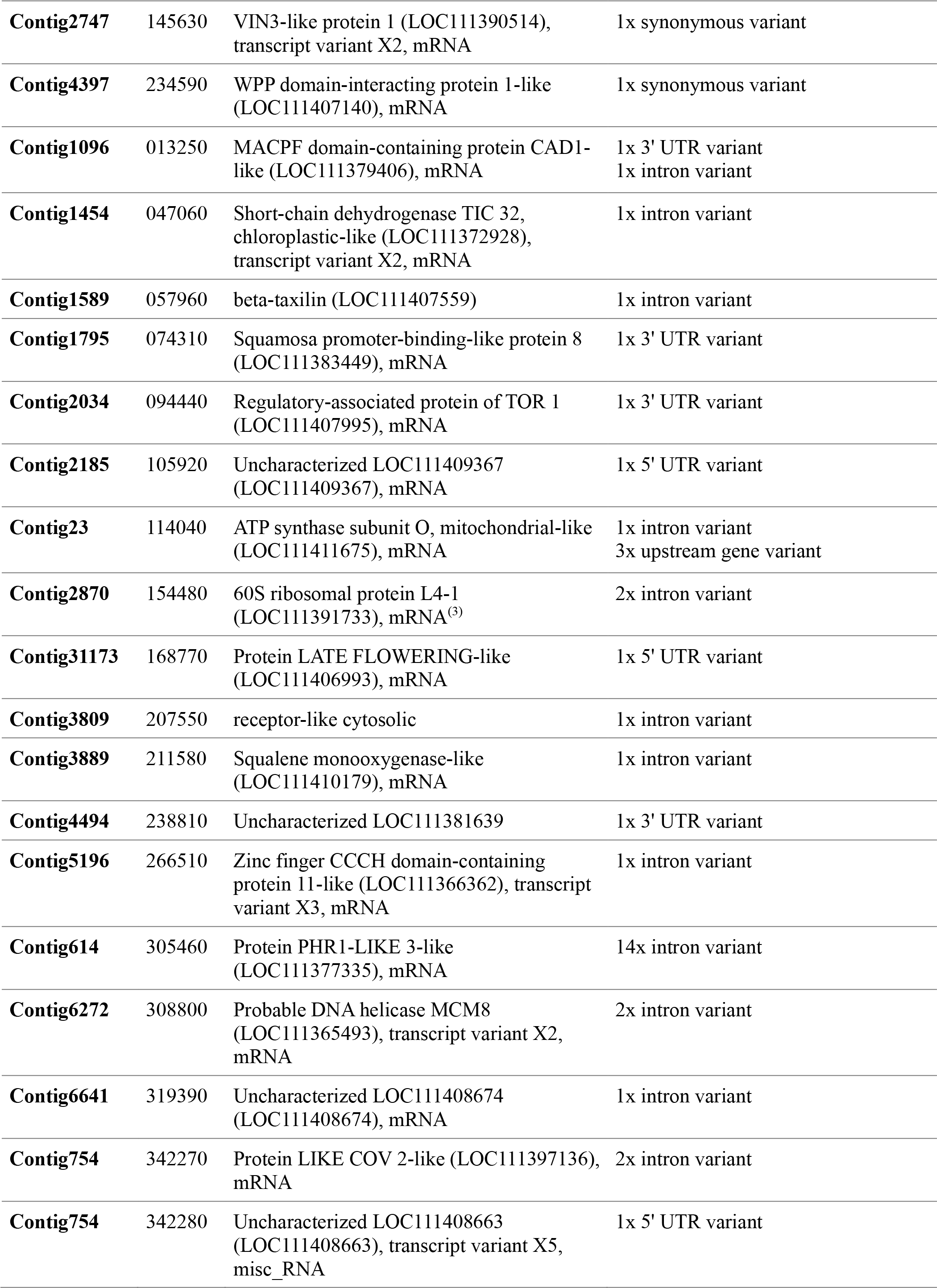

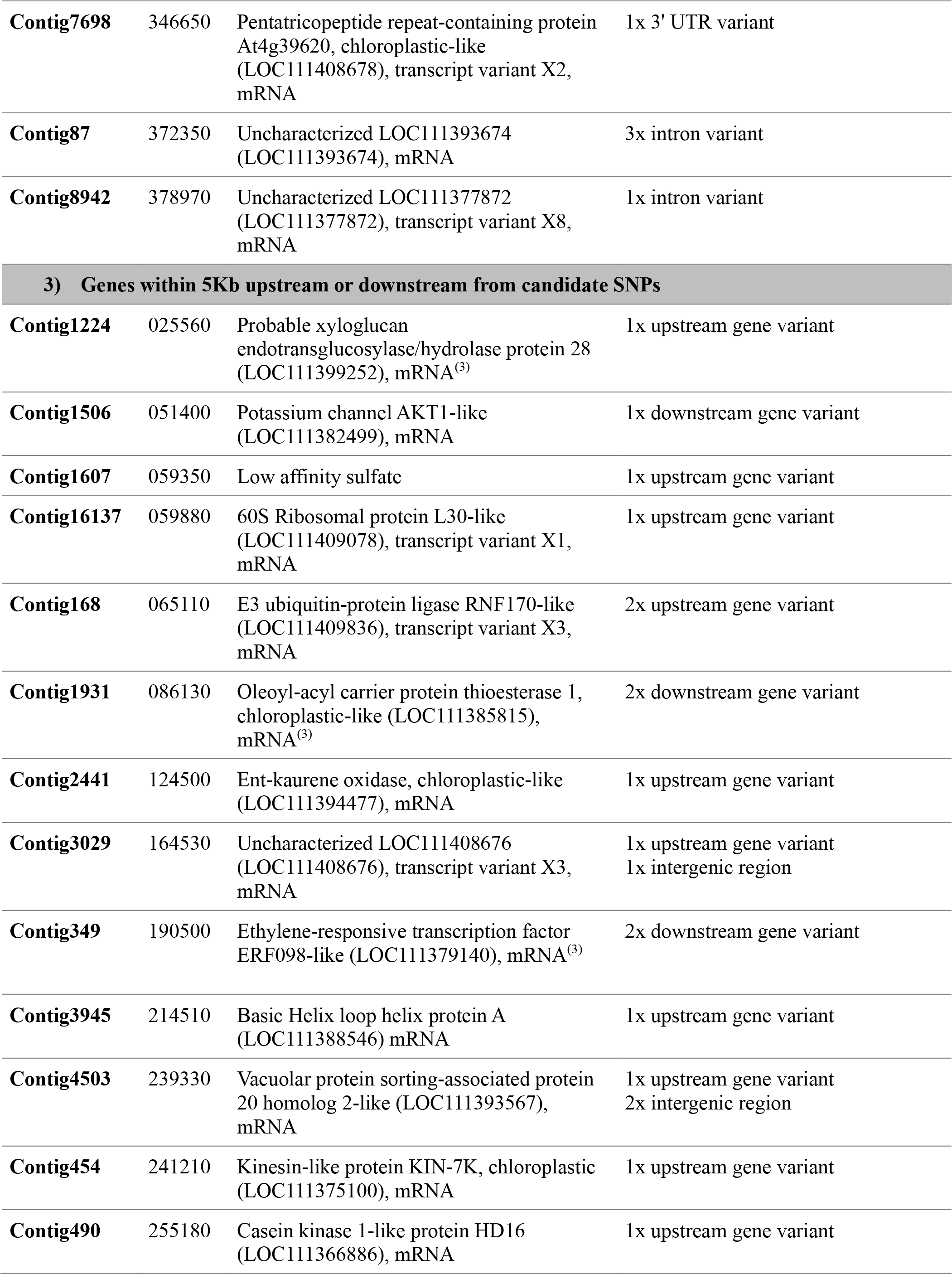

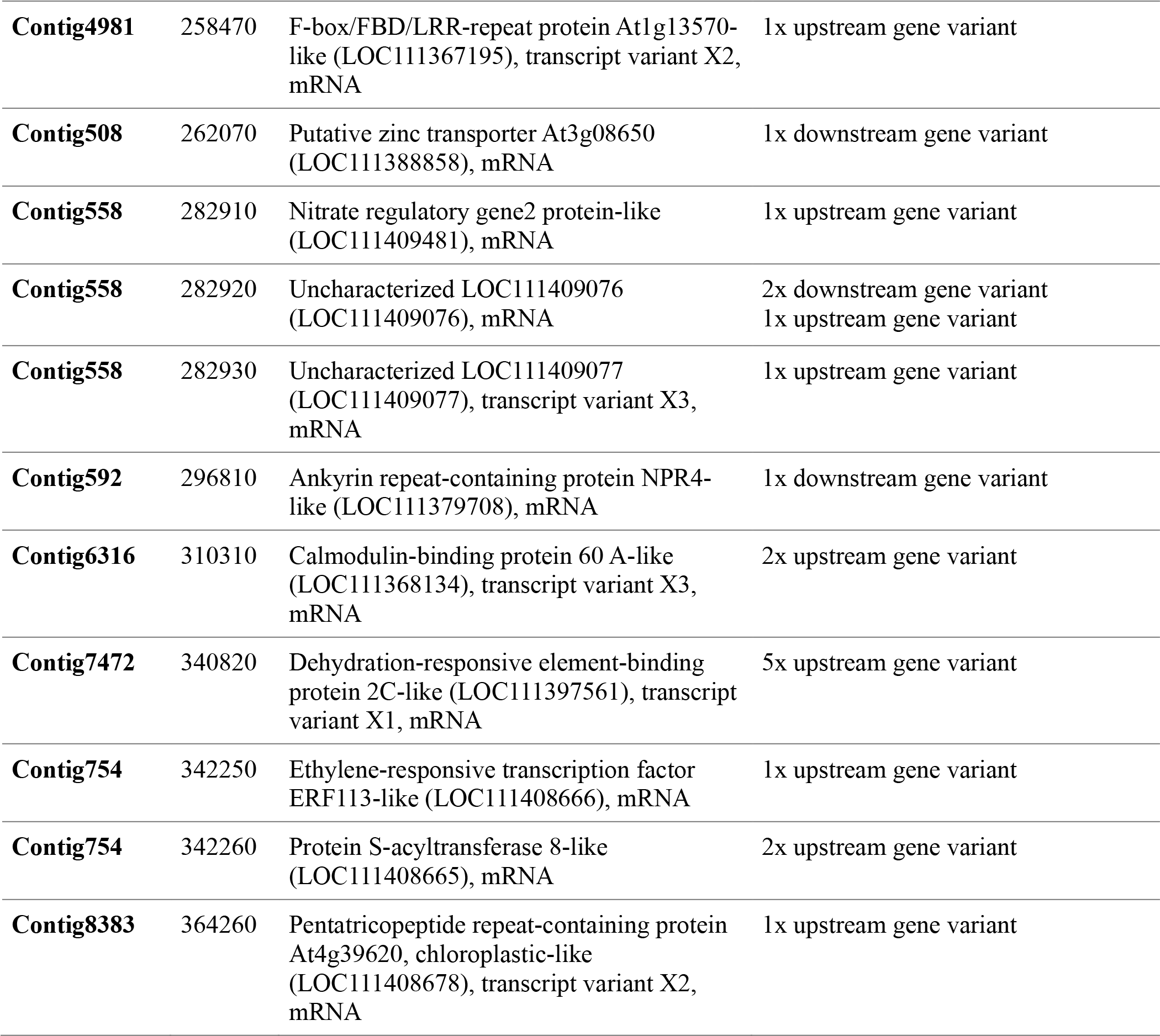

**Figure 3.**
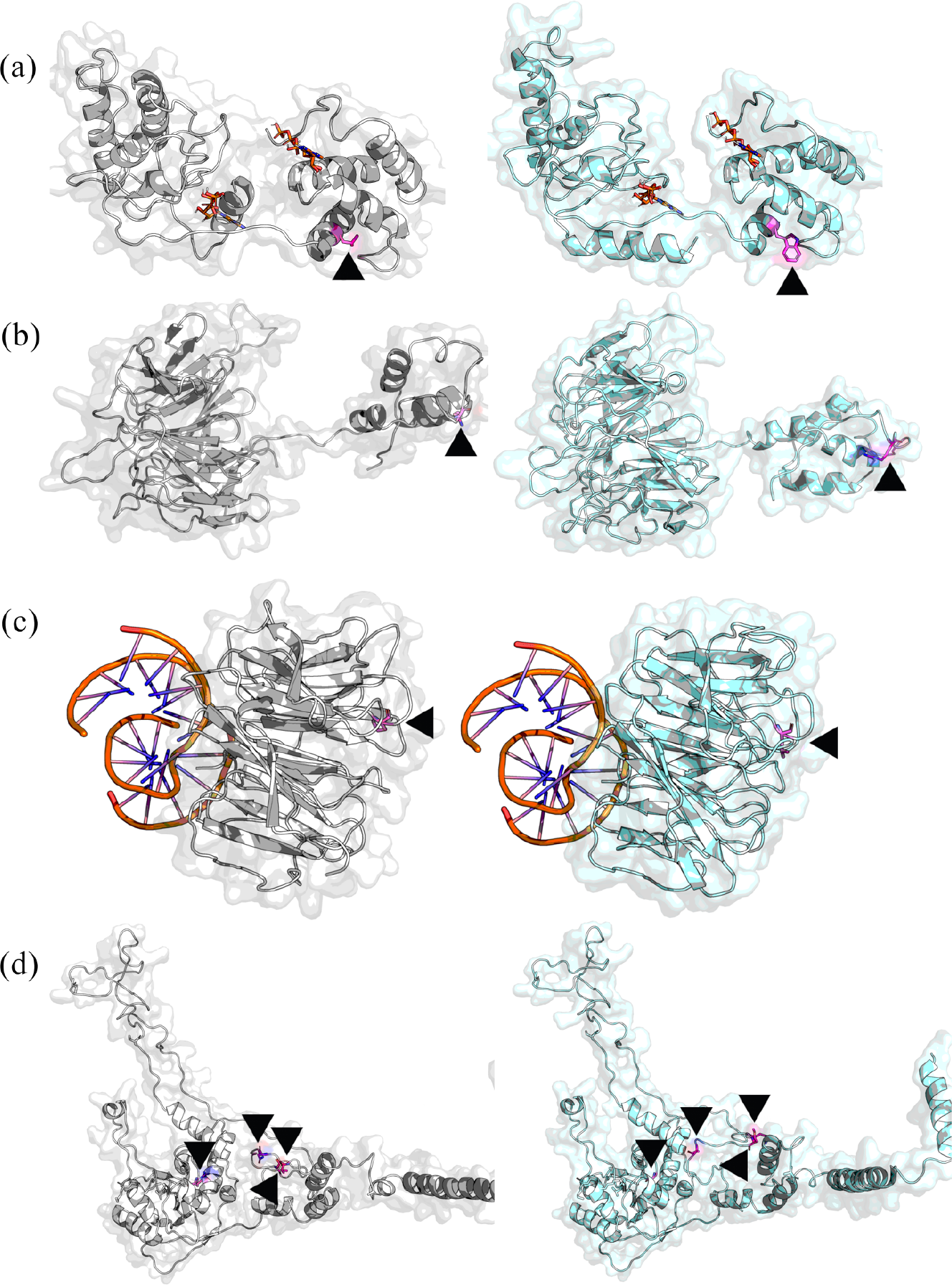
Predicted protein structures for genes containing amino acid changes associated with tree health status under ADB pressure. The protein structures to the left were more common in damaged trees, and those to the right were more common in healthy trees. Variant amino acids are coloured in magenta and indicated with a black arrowhead. (a) Gene FRAEX38873_v2_000003260, a BED finger-NBS-LRR resistance protein, where position 157 is a leucine (left) versus tryptophan (right) variant. Two ATP molecules are shown in orange to indicate the location of nucleotide binding sites. (b) Gene FRAEX38873_v2_000164520, a F-box/kelch-repeat, where position 13 is a glutamine (left) versus arginine (right) variant. (c) FRAEX38873_v2_000180950, a Protein DAMAGED DNA-BINDING, where position 99 is a proline (left) versus leucine (right) variant. DNA molecules are shown in orange docked at the proteins’ DNA binding sites. (d) Gene FRAEX38873_v2_000116110, a 60S ribosomal protein L4-1, where position 251 is an arginine (left) versus glycine (right) variant, position 285 is a methionine (left) versus arginine (right) variant, position 287 is an asparagine (left) versus lysine (right) variant and position 297 is a threonine (left) versus alanine (right) variant.

The other three genes with missense mutations have putative homologs with functions that have not been previously linked directly to disease resistance. Gene FRAEX38873_v2_000116110 is a 60S ribosomal protein L4-1 (RPL4-1) homolog, with four missense and nine synonymous variants associated with ADB damage level. The amino acid positions affected are in disordered regions in close proximity to one another (Figure 3d). Changes in this gene may affect the efficiency of mRNA translation^20^. Gene FRAEX38873_v2_000346660 is a Heat Intolerant 4 like protein with a phenylalanine to leucine variant. Gene FRAEX38873_v2_000180950 is a homolog of Damaged DNA-Binding 2 (DBB2), which has a role in DNA repair^21^ and contains a proline/leucine substitution within its WD40 protein binding domain (Figure 3c). This gene is found on Contig 332 between two G-type lectin S-receptor-like serine/threonine-protein kinase LECRK3 genes (FRAEX38873_v2_000180940 and FRAEX38873_v2_000180960) whose putative homologs are involved in brown planthopper resistance in rice^22^.

A further 24 genes contain significant (p < 1 × e^−13^) SNPs encoding variants that are transcribed but not translated (Table 1) Of these, four match genes that have been previously identified as involved in disease resistance in other species. Gene FRAEX38873_v2_000234590 encodes a WPP domain-interacting protein 1-like, and WPP domains have been linked to viral resistance in potato^23^. Gene FRAEX38873_v2_000305460 encodes a PHR1-LIKE 3-like protein which may play a role in immunity^24^ via the salicylic acid and jasmonic acid pathways^25^. Gene FRAEX38873_v2_000013250 encodes a Membrane Attack Complex and Perforin (MACPF) domain-containing Constitutively Activated cell Death (CAD) 1-like gene, which controls the hypersensitive response via salicylic acid dependent defence pathways^26^. FRAEX38873_v2_000211580 is a Squalene monooxygenase-like gene involved in the synthesis of phytosterols^27^ which have a role in plant immunity^28^.

Other genes involved in regulation were found to have significant (p < 1 × e^−13^) non-translated variants. FRAEX38873_v2_000266510 is a zinc finger CCCH domain-containing protein 11-like that is likely to be involved in regulation, perhaps of resistance mechanisms^29^. FRAEX38873_v2_000047060 is a short-chain dehydrogenase TIC 32, chloroplastic-like gene that is involved in the regulation of protein import^30^. FRAEX38873_v2_000074310 is putatively homologous to a squamosa promoter-binding (SBP)-like protein 8 that controls stress responses in *Arabidopsis*^31^. Two genes with non-coding variants seem to affect phenology: gene FRAEX38873_v2_000145630 encodes a Vernalisation Insensitive 3 (VIN3) like protein 1^32^ and gene FRAEX38873_v2_000168770 encodes a Late Flowering-like protein.

Interestingly, significant non-translated variants were also found in categories of genes that had unexpectedly shown significant missense variants. Another 60S ribosomal protein L4-1 gene, FRAEX38873_v2_000154480 (in addition to FRAEX38873_v2_000116110, which contains four missense variants) contains two intron variants associated with ADB damage. There are only three loci in the ash genome reference assembly matching the *Arabidopsis* 60S RPL4-1 (AT3G09630) gene. Another putative DNA repair gene was also hit (in addition to FRAEX38873_v2_000180950, which had a missense variant); gene FRAEX38873_v2_000308800 encoding a probable DNA helicase MiniChromosome Maintenance (MCM) 8 protein.

Six genes with putative roles in disease resistance have significant (p < 1 × e^−13^) SNPs within 5Kb up- or down-stream of them and are the closest known genes to those SNPs (Table 1). FRAEX38873_v2_000296810 matches an ankyrin repeat-containing protein NPR4-like gene; in *Arabidopsis* the *NPR4* gene is involved in defence against fungal pathogens and in mediation of the salicylic acid and jasmonic acid/ethylene-activated signalling pathways^33^. FRAEX38873_v2_000190500 is a putative ethylene-responsive transcription factor ERF098-like gene which may be involved in regulation of disease resistance pathways^34^. Gene FRAEX38873_v2_000342260 is a palmitoyltransferase or protein S-acyltransferases (PATs) 8-like gene^35^, which is likely to have a role in protein trafficking and signalling; in *Arabidopsis*, some PATs regulate senescence via the salicylic acid pathway^36^. FRAEX38873_v2_000025560 encodes a probable xyloglucan endotransglucosylase/hydrolase protein 27 which may play a role in extracellular defence against pathogens^37,38^. FRAEX38873_v2_0000258470 encodes an F-box/FBD/LRR-repeat protein likely to be involved in ubiquitination (see above). FRAEX38873_v2_0000340820 is a putative dehydration-responsive element-binding protein 2C-like (DREB2C) gene which has a role in osmotic-stress signal transduction pathways^39^.

The closest genes to 49 of the 203 most significant GWAS SNPs (p < 1 × e^−13^) were between 5Kb and 100Kb distant (Table S4). These included some with previous evidence of disease resistance functions. Gene FRAEX38873_v2_000086110 is a Leucine-rich repeat receptor-like serine/threonine-protein kinase β-amylase (BAM) 3, which is involved in fungal resistance in *Arabidopsis*^40^. Gene FRAEX38873_v2_000291580 is a bHLH162-like transcription factor whose putative *Arabidopsis* homolog is induced by infection with the downy mildew pathogen *Hyaloperonospora arabidopsidis*^41^. Gene FRAEX38873_v2_000169770 is likely to be involved in vacuolar protein sorting which can play a role in defence responses^42^. A cluster of SNPs on contig1355 are located at approximately 13-kb from gene FRAEX38873_v2_000037990, a small ubiquitin-like modifier (SUMO) conjugating enzyme UBC9-like gene. Inhibition of SUMO conjugation in *Arabidopsis* causes increased susceptibility to fungal pathogens^43^. Gene FRAEX38873_v2_000282910 is a nitrate regulatory gene 2 (NRG2) which could mediate nitrate signalling or mobilisation in response to pathogens^44^. Gene FRAEX38873_v2_000340830 is a trichome birefringence-like (TBL) 33 gene; mutants of TBL genes in rice plants confer reduced resistance to rice blight disease^45^.

### Genomic prediction

From 150 individual trees sampled from NSZ 204 (Dataset B) we generated a total of 2.9Tbp in 19.5 billion reads. Each individual tree was sequenced to 22X genome coverage on average. Quality metrics and GC content were very similar to Dataset A (Table S1). On average the percentage of reads mapped to the reference genome assembly per sample was 98.4% and 32,443,401 SNPs were found with read depth > 9 and mapping quality > 15.

To evaluate the genomic estimated breeding values of ADB damage (GEBV), we used the pool-seq data as a training population and the 150 NSZ 204 individuals as a test population. We obtained highest accuracy (correlation of observed scores and GEBV, *r* = 0.37; frequency of correct allocations, *f* = 0.68) using the top 10,000 SNPs by p-value from the GWAS, of which 9,620 SNPs had been successfully called in the test population (Figure 4). Smaller and larger SNP-dataset sizes performed less well. With a view to using a subset of these SNP for prediction, we reran the analysis using a subset of the 25% with the largest (absolute) estimated effect sizes and found minimal effect on the correlation (Figure 4), again finding the best result with (25% of) the dataset of 10,000 SNPs. Estimated effect sizes for all SNPs with models trained on 100 to 50,000 SNPs are shown in Supplementary File 1.

**Figure 4.**
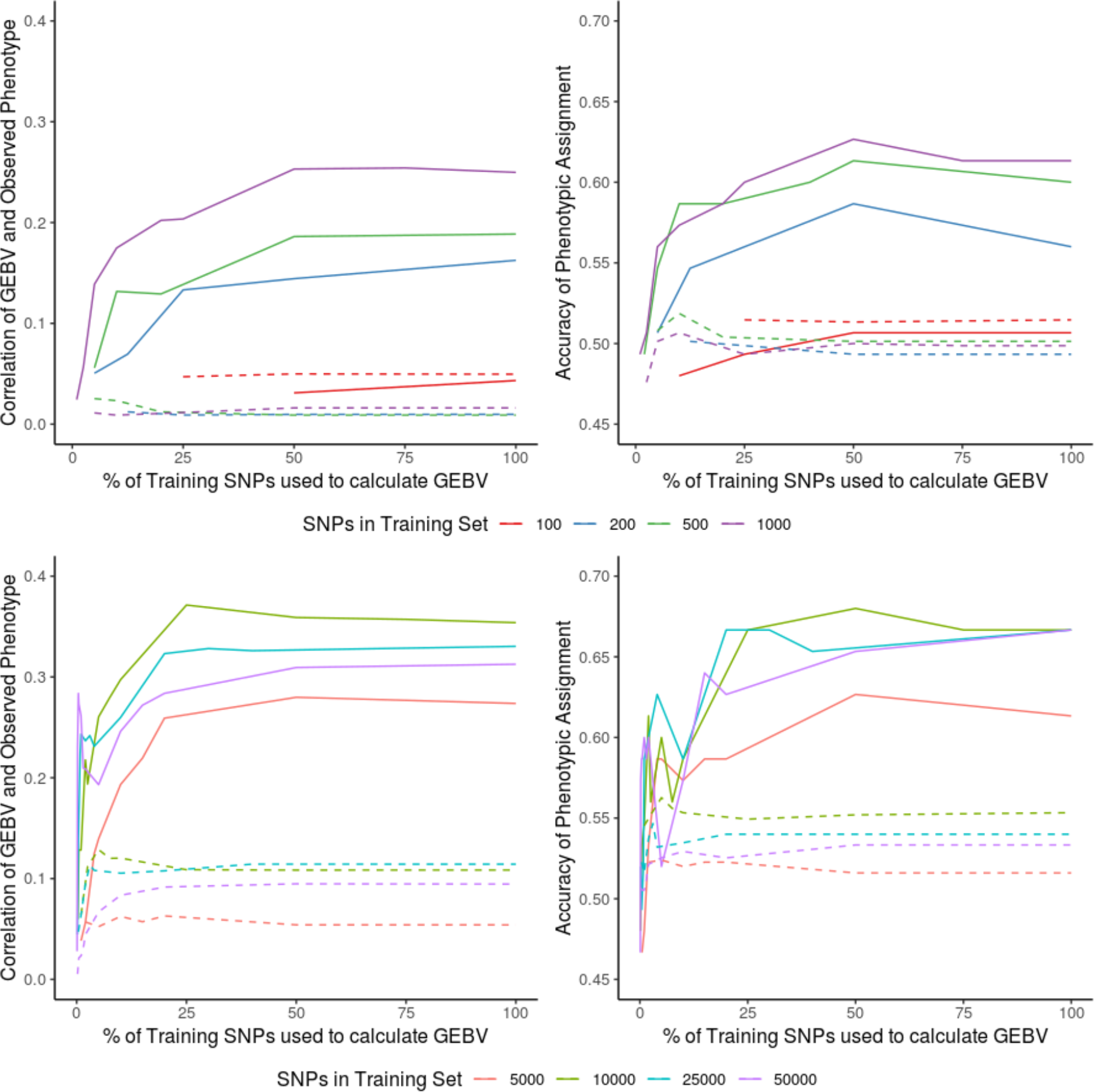
Genomic prediction of health under ash dieback pressure for 150 individual ash trees, with models trained on pooled sequencing of 1250 trees, using varying numbers of SNPs in training and test sets. Solid lines show results for SNPs selected using the pool-seq GWAS; dashed lines show average results using randomly selected SNPs. Left column: correlation of genomic estimated breeding value (GEBV) with observed health status. Right column: accuracy of health status assignment from GEBV.

Using the GWAS p-values as the criterion for selecting candidate SNPs for GP was far more effective than using a random selection from the genome, as judged by *r* and *f* scores (Figure 4). Despite this effect, there was not a strong association between the GWAS p-values and the effect size estimated by the genomic prediction: only 54 of the 2500 SNPs with the largest effect size were in the top 203 SNPs identified by the GWAS.

In a relatively small population with large heritable effects, spurious associations between some SNP alleles and a trait can arise. A sufficiently large number of randomly chosen SNPs will convey all the information on the relatedness of the individuals which, in turn, can be used to predict a trait simply because related individuals have similar trait values. To evaluate this effect, the 150 NSZ 204 individuals were used for GP as both a training dataset and a test dataset. The accuracy of the prediction with the top 50,000 GWAS-identified SNPs was no better than a random selection of 50,000 SNPs (Figure S5). Given this, we re-ran GP training on the pool-seq data with the pools from NSZ 204 (the seed source of the test population) excluded in case their inclusion had given spurious associations that contributed to the success of the first GP. This more stringent cross-validation showed a comparable performance to our previous GP trained on the full pool-seq dataset (maximum *r*= 0.36, *f*= 0.67; Figure S6).

For a breeding programme for increased resistance to ash dieback, accurate prediction of the most resistant trees is needed. We therefore examined the accuracy with which our highest GEBVs were assigning trees correctly to the undamaged health category. For the trees with the top 20% and 30% GEBV scores, we obtained predictive accuracies of *f*> 0.9, using as few as 200 predictive SNPs (Figure 5).

**Figure 5.**
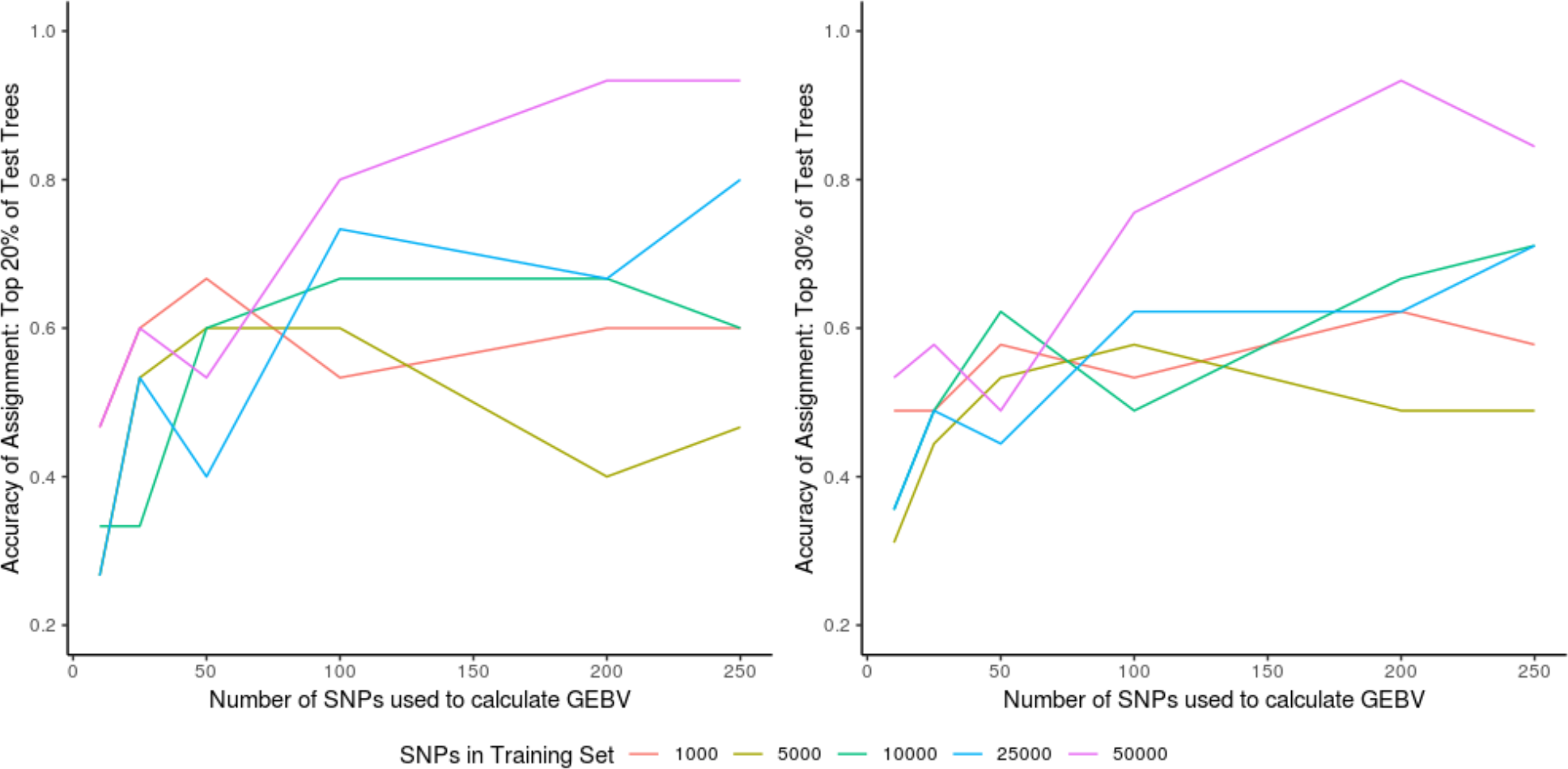
Genomic prediction accuracy of assignment of health status for the (left) top 20% and (right) top 30% of test population trees by GEBV, using 1000 to 50,000 SNPs identified by GWAS in the training set and use of ten to 250 SNPs in the testing set.

## Discussion

Many of the top SNP loci that we found associated with ash tree resistance to ash dieback are in, or close to, genes with putative homologs in other species that have been previously shown to detect pathogens, signal their presence, or regulate pathogen responses. Using SNPs identified by the GWAS to train GP on the pool-seq data, we obtained much greater accuracy in predicting the ADB damage score in 150 separate individuals than when we used the same number of randomly selected SNPs. Together, these results demonstrate we can use genotype to predict performance across different seed-sources, and that other genes that have not previously been implicated in plant pathogen resistance, such as 60S ribosomal protein L4-1 genes and some DNA repair genes, may be involved in resistance to ADB. None of our most significant SNPs were in or close to genes previously identified as showing gene expression changes associated with ADB resistance^3^, but we cannot exclude the possibility that our candidate SNPs may be controlling expression differences in these genes. The distribution of effect sizes and the predictivity peak using 2500 SNPs suggests that *F. excelsior* resistance to *H. fraxineus* is a highly polygenic trait and may therefore respond well to artificial and natural selection, allowing the breeding or evolution of durable increased resistance.

The levels of accuracy which our GP reached are high, and comparable to those that are used to inform selections in crop^46–50^, tree^12,51^ and livestock breeding programmes^52,53^. Thus, our results have the potential to increase the speed at which we can successfully breed ash dieback resistant trees. A common short-coming of GP is that predictions are highly population specific^12,54,55^, and the success of GP using randomly selected SNPs when training models within the individually sequenced trees suggests that population-specific GP can be easily made for ash. However, we made successful predictions in the individually sequenced trees using the pool-seq trained GP even when the pool-seq data for their seed-source provenance was not used in training the model. This suggests we have successfully identified widespread alleles that are involved in ADB resistance in many populations. There may well be further population-specific alleles that our methods have not detected. This study is the first that we are aware of to use pool-seq data to train a trans-populational GP model. The success of this approach in European ash – a genetically variable species – suggests it may be useful in many other ecologically important species as a cost-effective approach to successful genomic prediction of evolving traits.

## Methods

### Trial design

This study is based on a Forest Research mass screening trial planted in spring 2013, comprising 48 hectares of trials on 14 sites in southeast England as described in Stocks et al. 2017^15^. Briefly, each site was planted with trees grown from seed sourced from up to 15 different provenances. These were 10 British native seed zones (NSZ 106, NSZ 107, NSZ 109, NSZ 201, NSZ 204, NSZ 302, NSZ 303, NSZ 304, NSZ 403, NSZ 405), Germany (DEU), France (FRA), Ireland (CLARE and IRL DON), and a Breeding Seedling Orchard (BSO) planted by Future Trees Trust (FTT) comprised of half-sibling families from “plus” trees across Britain.

### Phenotyping and sampling

In July/August 2017 fresh leaves for DNA extraction were sampled from four of the trial sites that had heavy ash dieback damage: sites 16 (near Norwich, Norfolk), 21 (near Maidstone, Kent), 23 (near Norwich, Norfolk) and 35 (near Tunbridge Wells, Kent). We selected healthy trees (scores 7 on the scale of Pliura *et al.* ^56^) and trees with considerable ash dieback damage (scores 4 and 5 on the scale of Pliura *et al.* ^56^). Initially a total of 1536 trees were sampled. Of these 623 healthy and 627 unhealthy trees were selected for pooled sequencing with the total number of trees for each seed source and health status described in Table S2 and Figure S1. For individual sequencing, we selected a further 75 healthy and 75 unhealthy trees from NSZ 204 that were not included in the pools from this seed source.

### DNA extraction and sequencing

Leaf samples were transported to the lab using cool boxes. Fresh Genomic DNA was extracted from liquid nitrogen frozen leaf tissue using the DNeasy Plant Mini Kit or the DNeasy 96 Plant Kit (Qiagen) and eluted in 70 μl of Qiagen AE buffer. Quantification of genomic DNA was performed using the Quantus™ Fluorometer on all extractions. DNA purity quality checks were carried out using the Thermo Scientific™ NanoDrop 2000 for nucleic acid 260/280 and 260/230 absorbance ratios. Of the total number of extractions, 1400 were selected based on DNA quantity and quality thresholds. A minimum concentration of >20 ng/μl, OD260/280 >1.7 and total amount >1.0 μg of DNA was necessary for the sample to pass. Of the 1400 samples, 1250 were separated for the pooling and sequencing procedures and will be referred to as dataset A. A separate 150 individuals from NSZ 204, that were not included in the pools, were selected for individual genotyping and will be referred to as dataset B.

For the pooling procedure equal amounts of DNA from each sample were pooled together based on their initial DNA concentrations, adjusting the total volume of each sample accordingly. Pooling was based on seed source origin and health status with two pools for each seed source, one healthy and the other damaged. A total of 31 pools were created (Figure S1), one being a technical replicate of the healthy trees from NSZ 204 that was made by independently repeating all quantification, quality and pooling steps on the same 40 trees. NSZ 106 and NSZ 107 had 4 pools each as the samples were divided to maintain an average of 42 trees per pool. These therefore provide biological replicates. Studies have shown that pools sizes as small as 12 have provided robust and reliable population allele frequency estimates^14,57^.

TruSeq DNA PCR-Free (Illumina) sequencing libraries were prepared, using 350 base pair inserts. All sequencing was carried out using HiSeq X at Macrogen (South Korea) with 150 paired end reads with the goal of achieving a whole genome coverage (based on the estimated genome size of the *F. excelsior* reference individual^3^ of 80x per pool (2x coverage per individual) for dataset A and 20x for dataset B.

### Mapping to reference and filtering

Trimmomatic v0.38 was used for read trimming and adapter removal. Leading and trailing low quality or N bases below a quality of 3 were removed. Reads were scanned with a 4-base wide sliding window, cutting when the average quality per base dropped below 15 and excluding reads below 36 bases long^58^. Reads were then aligned to the reference genome for *Fraxinus excelsior*, assembly version BATG0.5, using the Burrows-Wheeler Alignment Tool (BWA MEM)^59^, version 0.7.17 with default settings. The mapped reads were filtered for a mapping quality of 20 with samtools (v1.9). On average the percentage of reads mapped to the reference was 98.3% for dataset A and 98.4% for dataset B. For both datasets Sequence Alignment Map (SAM) and binary version (BAM) files were created using Samtools. Indels were detected and removed using Popoolation2^60^ scripts (identify-indel-regions.pl and filter-sync-by-gtf.pl) that include five flanking nucleotides on both sides of an indel. The position of repeats in the reference genome was annotated previously^3^ using RepeatMasker v. 4.0.5 (with option -nolow) and that information used to remove repeats from these data using the same removal script provided by Popoolation2.

### Genetic structure of provenances

Major allele frequency information was extracted from dataset A for each of the 31 populations using a modified output of the allele frequency differences script (snp-frequency-diff.pl) from the PoPoolation2 package. This table of major allele frequencies was imported and converted to a genpop object and subsequently analysed using the R package adegenet^61^. A Correspondence Analysis on genpop objects was performed in order to seek a typology of populations. Correlation between populations was calculated and plotted, for the major allele frequencies from dataset A, using the corrplot R package^62^.

### Genome wide association study

For dataset A the software package PoPoolation2^60^ was used to identify significant differences between damaged and healthy trees. For this an mpileup input was generated using Samtools followed by the creation of a file that had all the variants synchronized across the pools and requiring a base quality of at least 20. The statistical test to detect allele frequency changes in biological replicates was the Cochran-Mantel-Haenszel (CMH) test^63^. With this test a 2×2 data table was created for each seed source (15) with two phenotypes (healthy and damaged) and the two major alleles for each SNP. The counts of each allele for each phenotype were treated as the dependent variables. The parameters set for PoPoolation2, given there were 30 pools with DNA from 1250 individuals, were: min count 15 (minimum allele count to be included), min coverage 40, max coverage 3000. False discovery rate control was performed using the R package q-value^64^. We excluded contig 18264 from the reference sequence because it appears to be derived from fungal contamination: its top BLAST hit in the GenBank nucleotide collection is to nrDNA in a species of the fungal genus *Phoma* (MH047199.1), a putative fungal endophyte.

Putative functions for genes containing, or near, the pool-seq GWAS top SNPs were assigned by obtaining the CDSs from the Ash Genome website^3^ and using the command line NCBI Basic Local Alignment Search Tool (BLAST+) optimized for the megablast algorithm to search the GenBank Nucleotide database. The top result for every BLAST search was extracted and their predicted gene functions were used to functionally annotate the ash genes. Any search that yielded no matches when using megablast was then repeated using the blastn algorithm and ultimately cDNA sequences if the latter was also uninformative. Potential functional impacts for each of the top 203 GWAS SNP loci were determined using SNPeff (v4.3T)^65^. A custom genome database was built from the *F. excelsior* reference assembly using the SnpEff command “build” with option “−gtf22”; a gtf file containing the annotation for all genes, as well as fasta files containing the genome assembly, CDS and protein sequences, were used as input. Annotation of the impact of the 203 SNPs was performed by running SnpEff on all *F. excelsior* genes with default parameter settings.

### Protein modelling

Proteins containing SNPs identified by SnpEff as coding for amino acid substitutions were modelled. Protein coding sequences were taken from the predicted proteome of the BATG 0.5 reference genome^3^ and modelled both with the amino acid(s) associated with ADB damage in our GWAS, and with the amino acid(s) associated with healthy trees. Models were predicted using three methods: RaptorX-Binding (http://raptorx.uchicago.edu/BindingSite/), Swiss-modeller^66^ and Phyre2^67^. These models were compared by manually alignment in PyMOL v.2.0^68^, and only those with congruent models were taken forward, based on their Phyre2 and RaptorX-Binding models. Potential binding sites and candidate ligands were analysed using RaptorX-Binding and literature searches. SDF files for candidate ligands were obtained from PubChem (https://pubchem.ncbi.nlm.nih.gov) and converted to 3d pdb files using Online SMILES Translator and Structure File Generator (https://cactus.nci.nih.gov/translate/). Docking with our protein models was analysed using Autodock Vina v.1.1.2^69^ with the GUI PyRx v.0.8^70^. Following docking, ligand binding site coordinates were exported as SDF files from Pyrex and loaded into PyMOL with the corresponding protein model file for the “healthy” and “damaged” protein models. Binding sites were then annotated and the variable residues were labelled. Possible RNA and DNA binding sites were predicted using DRONA (http://crdd.osdd.net/raghava/drona/links.php). The presence of signal peptides were detected using SignalP 4.1 server and Phobius server (http://phobius.sbc.su.se/index.html); both were run with default parameters and for Phobius the “normal prediction” method was used. The presence of a signal peptide was confirmed only if it was predicted by both methods. Motif search (https://www.genome.jp/tools/motif/) and ScanProsite (https://prosite.expasy.org/scanprosite/) were used to predict protein domains and their locations for our candidate genes.

### Genomic Prediction

We trained a GP model based on the pool-seq data (Dataset A). Subsets of 100, 200, 500, 1000, 5000, 10000, 25000 and 50000 SNPs with the most significant GWAS results were selected from Dataset A and used as a training set. Results were compared with SNP sets of the same size drawn at random from the genome. SNPs from contig 18264 (suspected to be fungal contamination) were excluded. We constructed a pipeline available at https://github.research.its.qmul.ac.uk/btx330/gppool. The vector of ADB damage scores for each pool, y, was predicted by the rrBLUP model as: y = **X**β + ε, where β is a vector of allelic effects (treated as normally distributed random effects), and the residual variance is Var[ε]. The genetic data are encoded in the design matrix **X** which has a row for each pool and a column for each SNP allele. The entry for pool *p* and locus *l* is X[p,l] = *f*_*pl*_ − *μ*_*s*_, where *f*_*pl*_ is the frequency of the focal allele and *μ*_*s*_ is its mean frequency across the pools from the same seed-source as *p*.

The Reduced Maximum Likelihood solution to the model was obtained using the *mixed.solve* function in rrBLUP v4.6^71^ to give estimated effect sizes (EES) for the minor and major alleles at each SNP under consideration. Subsets of the 10 – 50,000 SNPs with the greatest EES were used to predict GEBV for each of the 150 individuals from provenance NSZ 204. For these individuals (dataset B) variant calling was performed using bcftools with the raw set of called SNPs filtered using VCFtools (vcfutils) - set at minimum read depth of 10 and minimum mapping quality 15. Filtering of loci was carried out using thresholds of >95% call rate and >5% MAF. Samples were filtered based on a >95% call rate and <1% inbreeding coefficient. SNPs were also filtered if they deviated significantly from Hardy-Weinberg equilibrium. GEBV was calculated as the sum the EES and the relative frequency of each focal allele. Predictions were repeated with seed-source NSZ 204 excluded from the training dataset to avoid spurious correlations due to population stratification.

Test trees were assigned to high and low susceptibility groups based on their GEBV and the accuracy of the assignment was tested using the formula: *f* = correct assignments/total assignments, with correct assignments defined as those that corresponded to the observed phenotypes. Correlation of GEBV and phenotypic classification, *r*, was calculated using the Pearson correlation coefficient.

We also carried out genomic prediction based solely on the 150 individuals in Dataset B. A ratio of 60/40 was used for training and testing populations and missing markers were imputed using the function R package A.mat^72^ with default settings. SNPs were selected from the GWAS output ordered by p-value. A total of 100, 500, 1000, 5000, 10000, 50000, 100000, 250000, 500000, 1000000 and 5000000 SNPs were selected from each filtered set and used for training and testing of the GP model. The same number of SNPs were selected at random (using R) from the fully filtered dataset and also used for training and testing the GP model. We used using the *mixed.solve* function in rrBLUP v4.6^71^ and Genomic Selection in R course scripts available at http://pbgworks.org. A total of 500 iterations were run of the rrBLUP. For the randomly selected SNPs, the 500 iterations were repeated ten times.

### Data and software availability

The authors confirm that all raw or analysed data supporting this study will be distributed promptly upon reasonable request. All trimmed reads are available at the European Nucleotide Archive with primary accession number: PRJEB31096. The gppool pipeline developed as part of the project to run GP trained on pool-seq data can be found at https://github.research.its.qmul.ac.uk/btx330/gppool. All software used (Trimmomatic, BWA, Samtools, Bcftools, VCFtools, PoPoolation2, R, Repeatmasker, SNPeff, Haploview, NCBI BLAST, RaptorX-Binding, Swiss-modeller, Phyre2, SMILES, Autodock Vina v.1.1.2, PyRx v.0.8, PyMOL, DRONA, SignalP 4.1 server, Phobius server, NetPhos 3.1 Server and Group-based Prediction System (GPS)) is commercially or freely available.

## Acknowledgements

This study was supported by Forest Research (FR), Queen Mary University of London (QMUL) and the Royal Botanic Gardens Kew. J.J.S. is funded by a Conselho Nacional de Pesquisa e Desenvolvimento (CNPq) studentship and is part of the Brazilian Scientific Mobility Program – Science without Borders (SwB). S.L. and R.J.A.B. were partly funded by Living with Environmental Change (LWEC) Tree Health and Plant Biosecurity Initiative - Phase 2 grant BB/L012162/1 funded jointly by the BBSRC, Defra, Economic and Social Research Council, Forestry Commission, NERC and the Scottish Government. FR designed and setup field trials. Funding for the field trails was supplied by the Department for Environment, Food and Rural Affairs (DEFRA) contract number TH032 ‘Rapid screening for Chalara resistance using ash trees currently in commercial nurseries’ with additional financial contribution from Department of Agriculture, Food and the Marine, Ireland, and donation of trial trees from Maelor Forest Nurseries. R. J. A. B. and L. J. K were also supported in this work by funding from the Defra Future Proofing Plant Health scheme and the Erica Waltraud Albrecht Endowment Fund. Sequencing was paid for by a direct grant from Defra to RBG Kew.

## Author Contributions

J.J.S. performed the field assessments and sampling, data analysis for all the GWASs, GS for dataset B and wrote the manuscript. R.J.A.B supervised field work, data analysis and interpretation and wrote the manuscript. L.J.K. analysed genetic data. S.J.L designed the field trials. R.A.N designed the statistical approaches. C.L.M developed and performed methods for Genomic Prediction with training on pool-seq data. W.P modelled the proteins. All authors reviewed the manuscript.

## Supplementary Information

**Supplementary Table 1.** Sequencing, Quality and Mapping values for each Dataset (A and B).

| Item | Dataset A (pool-seq) |  |  | Dataset B (individuals) |  |  |
| --- | --- | --- | --- | --- | --- | --- |
|  | Average | Min | Max | Average | Min | Max |
| Read Bases | 7.70E+10 | 7.24E+10 | 7.95E+10 | 1.95E+10 | 1.79E+10 | 1.99E+10 |
| Reads | 5.10E+08 | 4.79E+08 | 5.27E+08 | 1.29E+08 | 1.19E+08 | 1.32E+08 |
| GC(%) | 35.21 | 35.03 | 35.45 | 35.14 | 34.72 | 35.51 |
| AT(%) | 64.79 | 64.55 | 64.97 | 64.86 | 64.49 | 65.28 |
| Q20(%) | 96.44 | 94.13 | 97.51 | 97.20 | 96.57 | 97.86 |
| Q30(%) | 92.26 | 87.87 | 94.35 | 93.71 | 92.37 | 95.13 |
| Mapped (%) | 98.3 | 97.4 | 98.8 | 98.4 | 93.3 | 99.1 |

**Supplementary Table 2.** Distribution of samples in pooled dataset (A) and individually genotyped dataset (B) according to site and seed source.

| **Provenances** | **Site 16** | **Site 21** | **Site 23** | **Site 35** | **Total** |
| --- | --- | --- | --- | --- | --- |
| DEU | 28 | 79 |  |  | 107 |
| FTT SO | 9 | 32 |  | 34 | 75 |
| IRL DON | 38 | 10 |  | 26 | 74 |
| NSZ 106 | 54 | 62 | 12 | 60 | 188 |
| NSZ 107 | 20 | 80 | 18 | 50 | 168 |
| NSZ 109 | 10 | 50 | 12 | 16 | 88 |
| NSZ 201 | 11 | 55 | 4 | 10 | 80 |
| NSZ 204 | 60 | 11 | 4 | 6 | 81 |
| NSZ 302 | 14 | 32 | 14 | 39 | 99 |
| NSZ 303 | 6 | 20 | 8 | 36 | 70 |
| NSZ 304 | 14 | 38 | 8 | 17 | 77 |
| NSZ 403 | 18 | 18 | 14 | 32 | 82 |
| NSZ 405 | 1 | 17 | 10 | 33 | 61 |
| **Total** | **283** | **504** | **104** | **359** | **1250** |

*Individual Samples (Dataset B)*
| **Provenances** | **Site 16** | **Site 21** | **Site 23** | **Site 35** | **Total** |
| --- | --- | --- | --- | --- | --- |
| NSZ 204 | 58 | 51 | 17 | 24 | 150 |
| **Total** | **58** | **51** | **17** | **24** | **150** |

**Supplementary Table 3.** Comparison of the number of significant calls for the p-values, estimated q-values, and estimated local FDR values.

|  | **<1e-04** | **<0.001** | **<0.01** | **<0.025** | **<0.05** | **<0.1** | **<1** |
| --- | --- | --- | --- | --- | --- | --- | --- |
| p-value | 102,440 | 287,612 | 821,046 | 1,258,574 | 1,752,684 | 2,459,337 | 9,347,124 |
| q-value | 4,275 | 19,337 | 110,003 | 232,006 | 410,712 | 735,089 | 7,942,196 |
| local FDR | 3,149 | 10,395 | 57,370 | 121,222 | 213,502 | 379,746 | 3,360,672 |

**Supplementary Table 4.**
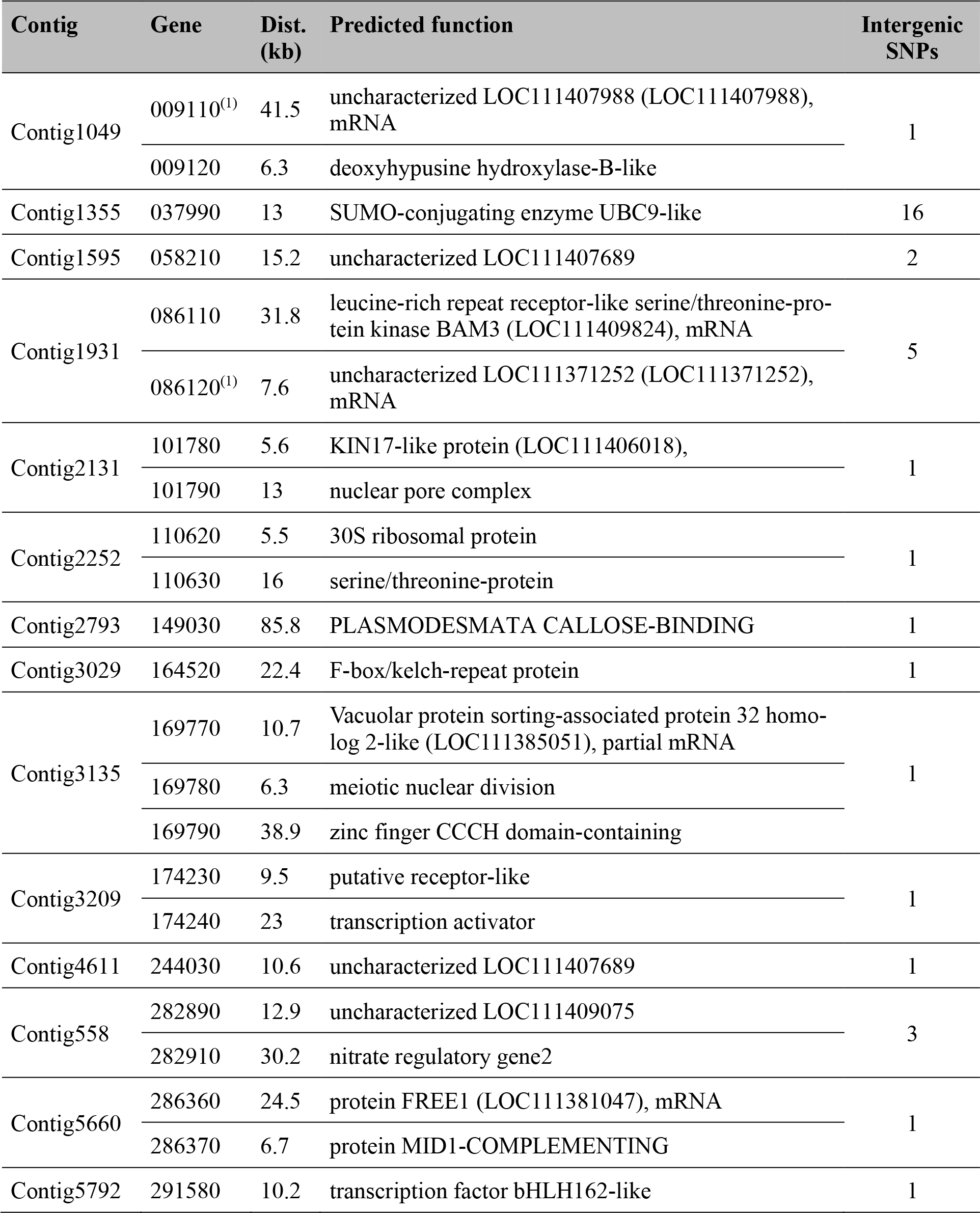
List of ash genes closest to the subset of the top 203 GWAS candidate SNPs (with −log_10_(p) > 13) that are over 5Kb from an annotated gene. Genes up to 100Kb from SNPs are shown. The “Gene” column gives the final six digits for the full gene names for the annotation of the ash genome^11^, which are in the form FRAEX38873_v2_000######. The column “Dist.” shows the distance of the gene from the nearest GWAS SNP. The predicted functions are from the olive genome.

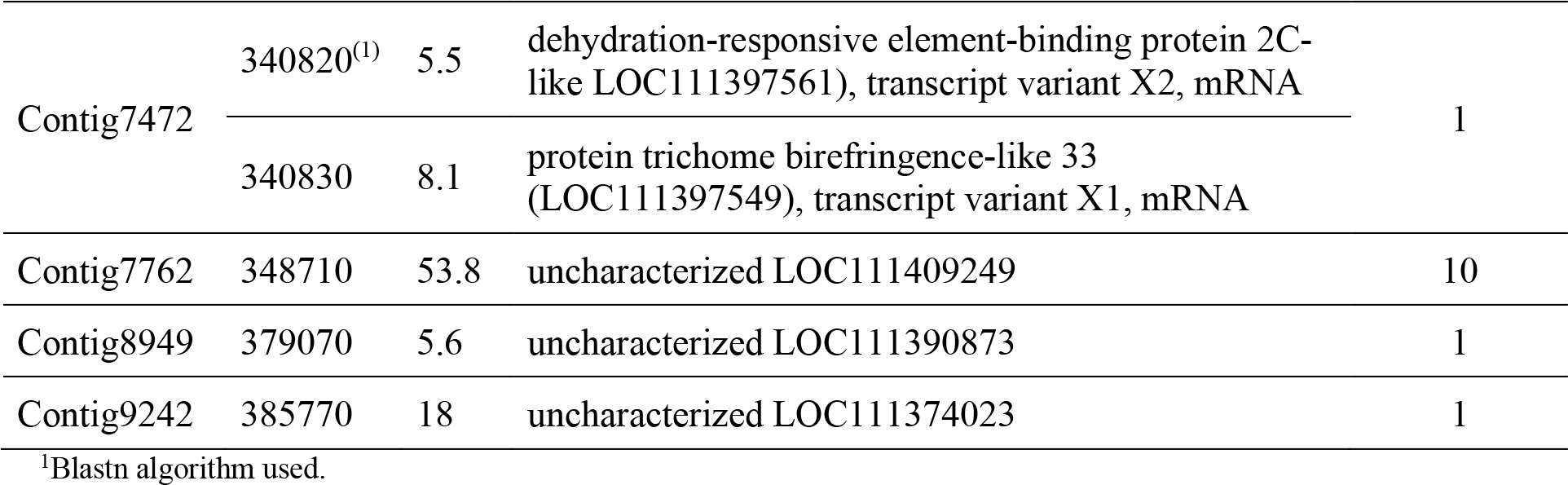

**Supplementary Table 5.** Polymorphic amino acid allele identities and frequencies at significant GWAS loci found in the top 203 hits by p-value (with −log_10_(p) > 13)

| Contig | Gene | Predicted function | Major allele | Minor allele | Position in protein | MAF in healthy trees | MAF in damaged trees |
| --- | --- | --- | --- | --- | --- | --- | --- |
| Contig10122 | 003260 | BED finger-NBS-LRR resistance protein | Leu | Trp | 157 | 0.216 | 0.121 |
| Contig10122 | 003270 | Protein CPR-5-like | Ile | Ser | 36 | 0.216 | 0.121 |
| Contig2324 | 116110 | 60S ribosomal protein L4-1 | Gly | Arg | 251 | 0.285 | 0.382 |
| Contig2324 | 116110 | 60S ribosomal protein L4-1 | Arg | Met | 285 | 0.263 | 0.354 |
| Contig2324 | 116110 | 60S ribosomal protein L4-1 | Lys | Asn | 287 | 0.322 | 0.431 |
| Contig2324 | 116110 | 60S ribosomal protein L4-1 | Ala | Thr | 294 | 0.301 | 0.393 |
| Contig3029 | 164520 | F-box/kelch-repeat protein SKIP6 | Gln | Arg | 13 | 0.136 | 0.052 |
| Contig332 | 180950 | Protein DAMAGED DNA-BINDING | Pro | Leu | 99 | 0.266 | 0.140 |
| Contig614 | 305440 | Uncharacterized | Gly | Asp | 1155 | 0.211 | 0.341 |
| Contig7698 | 346660 | Protein HEAT INTOLERANT 4-like | Phe | Leu | 12 | 0.123 | 0.064 |

**Supplementary Figure 1.**
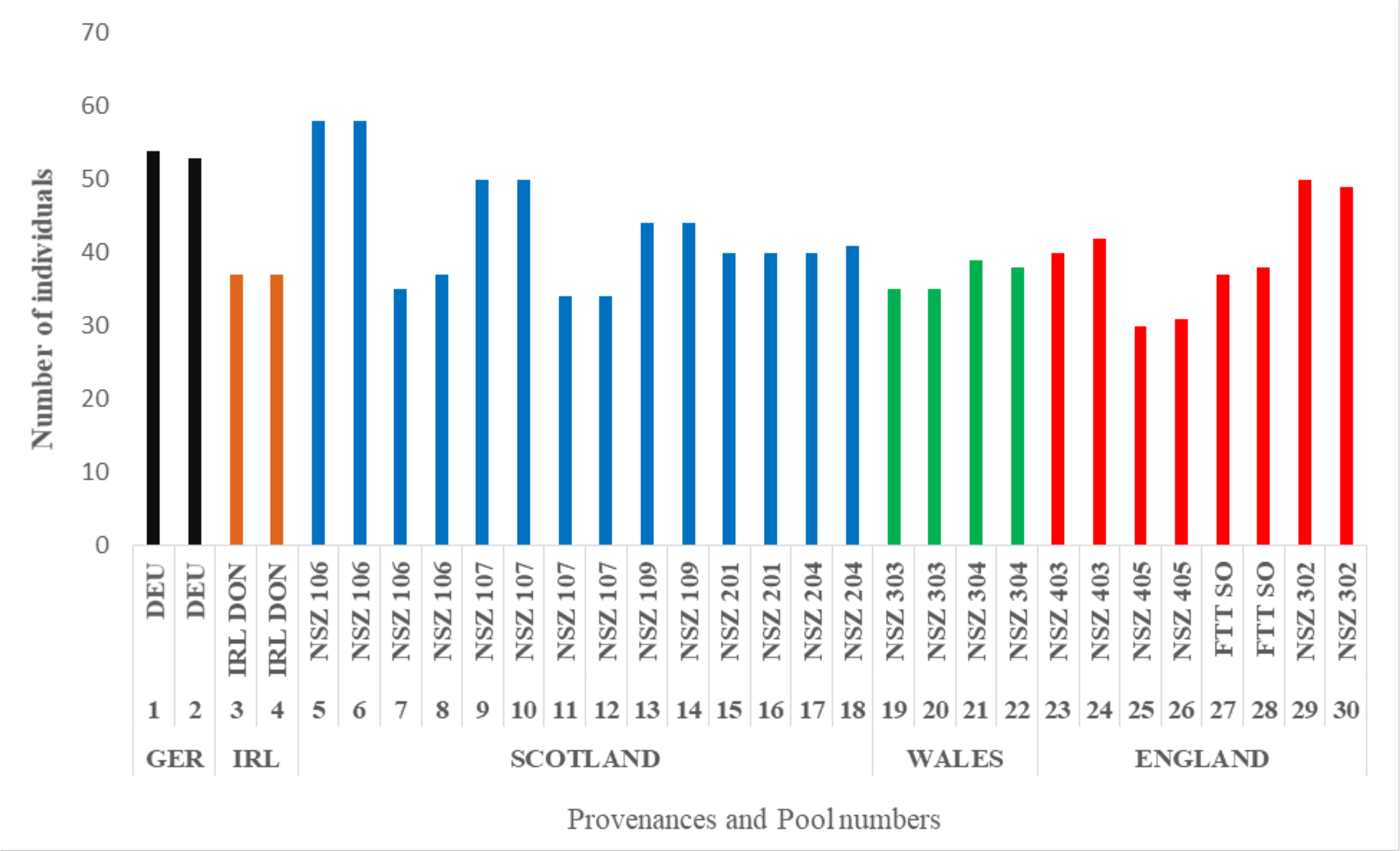
Number of individuals in each pool (odd pool numbers represent healthy and even numbers susceptible populations) and country of origin.

**Supplementary Figure 2.**
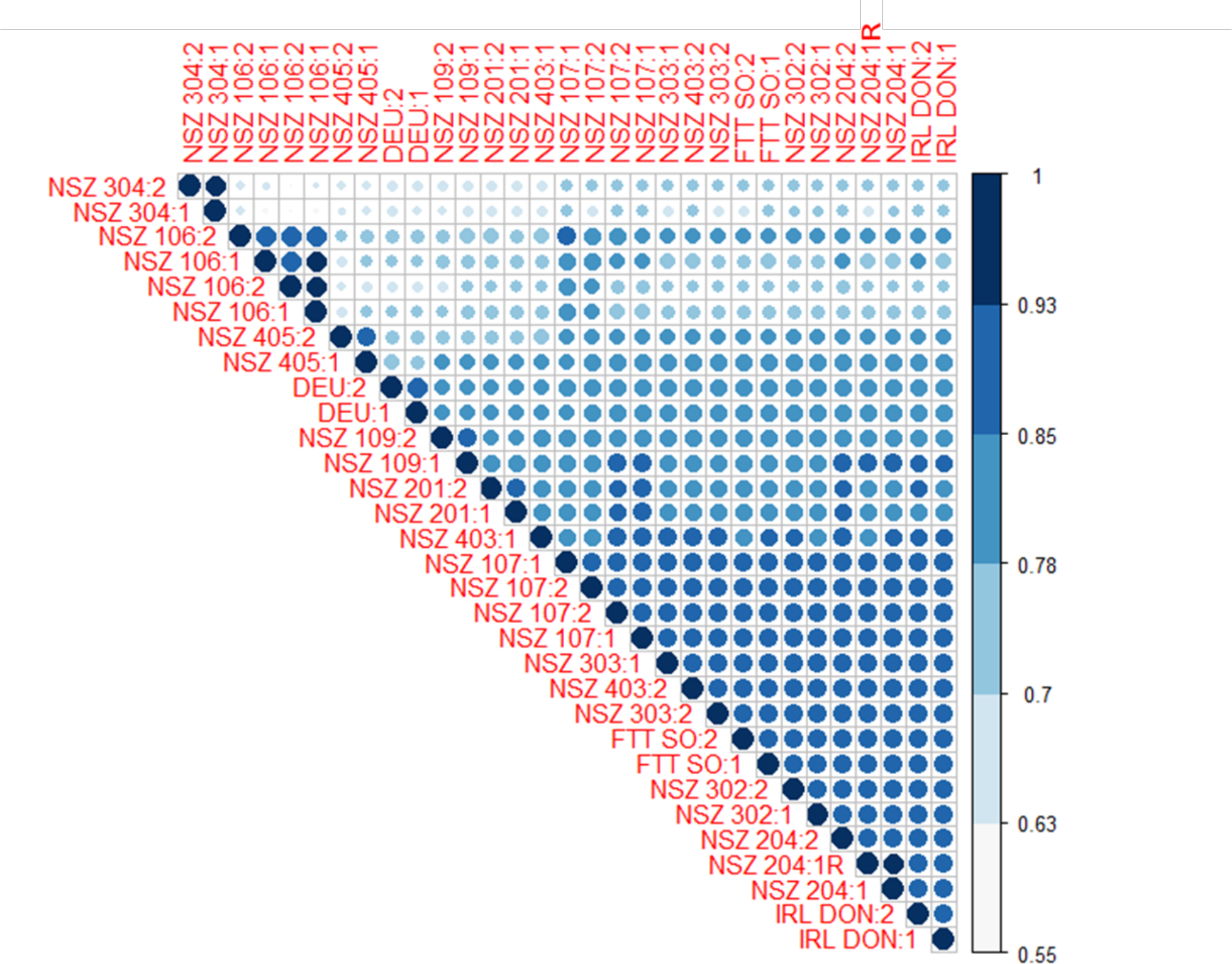
Circle plot of major allele frequency correlation values between all 31 pools. Numbers after seed source code correspond to health status (1 - healthy or 2 - damaged by ADB). Pool NSZ204:1 (with low ADB damage) was technically replicated (NSZ204:1R) using the same set of trees. Both pools from NSZ106 and NSZ107 were biologically replicated for both high and low damage pools, using different sets of trees.

**Supplementary Figure 3.**
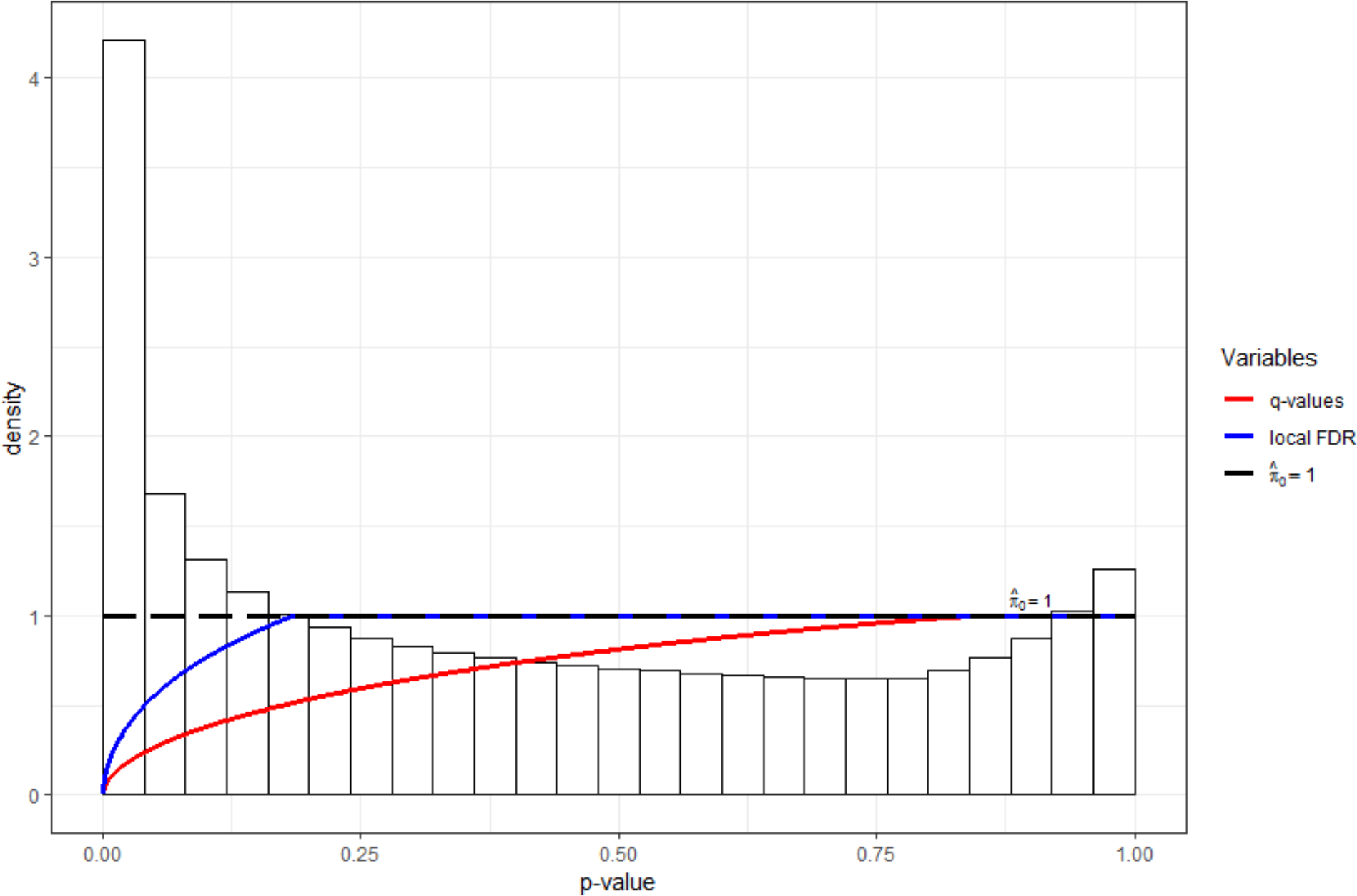
Pool-seq GWAS p-value density histogram with line plots of the q-values and local False Discovery Rate (FDR) values versus p-values. The π0 estimate is also displayed.

**Supplementary Figure 4.**
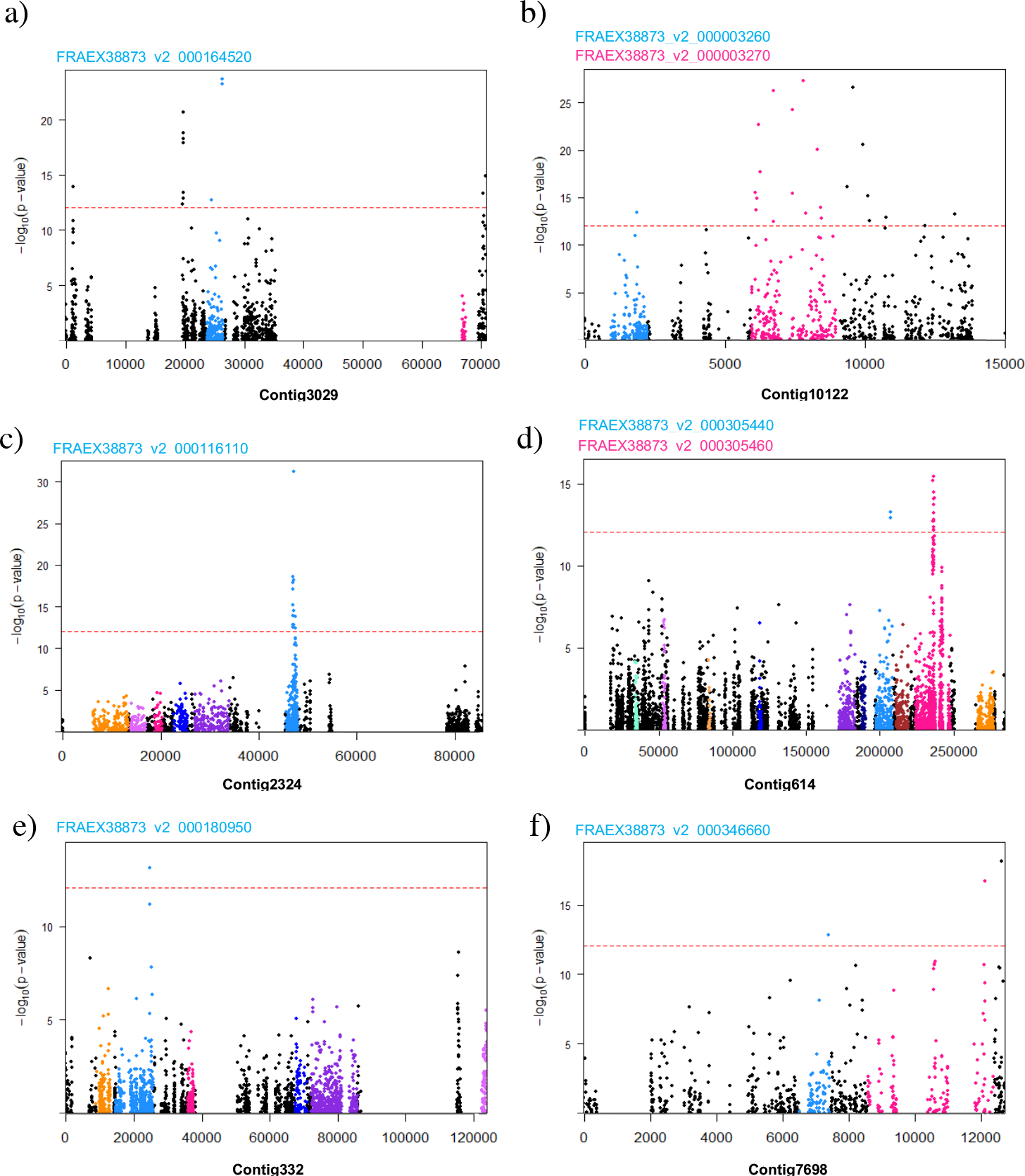
Manhattan plots for contigs containing genes in which SNPs encoding an amino acid substitution were in the top 203 pool-seq GWAS candidates. All genes present on the contigs are colored and those containing SNPs causing missense alterations to coding regions are labelled using the same colour as the gene’s SNPs in the Manhattan plot.

**Supplementary Figure 5.**
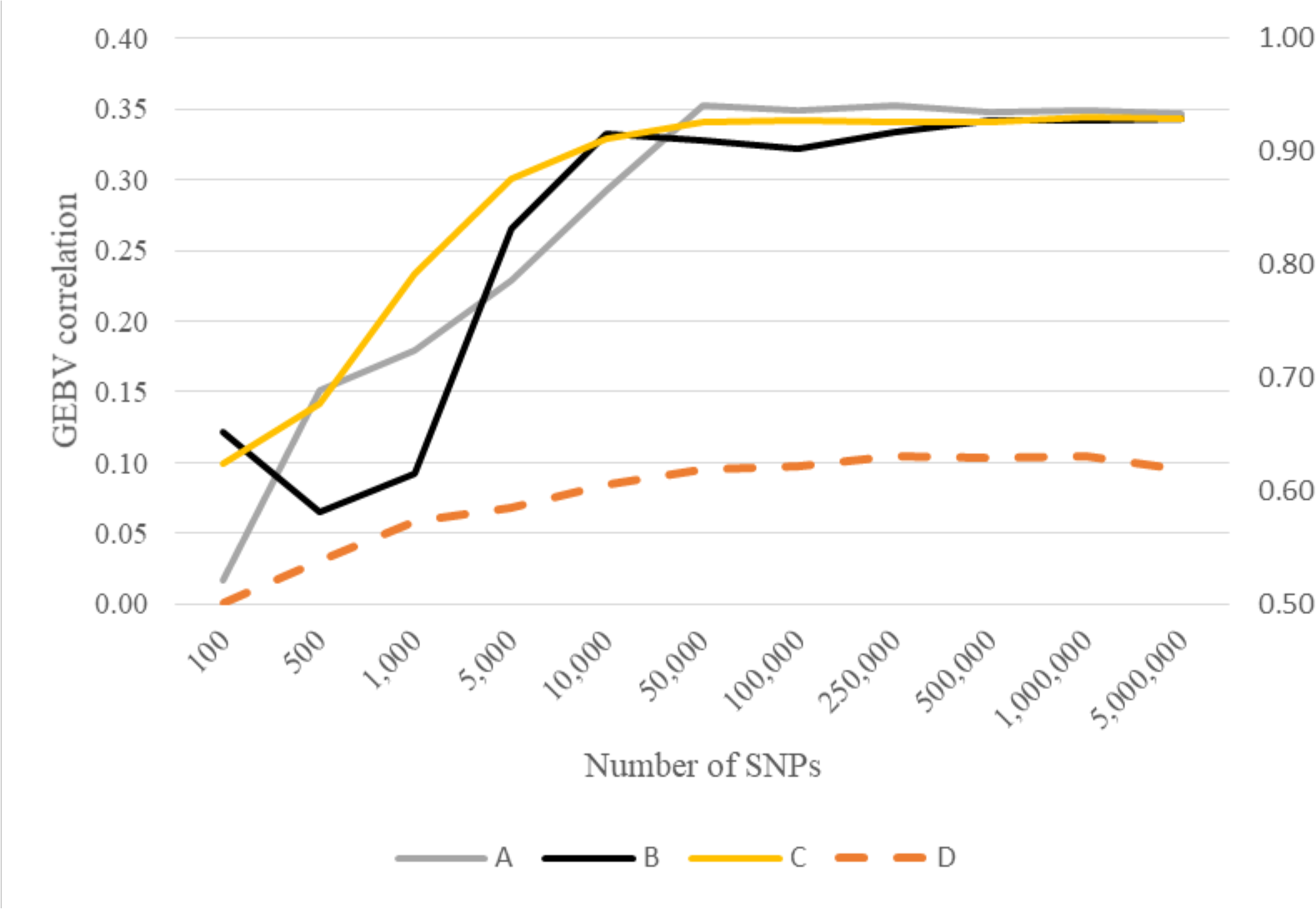
Genomic prediction results using the 150 individually genotyped samples as both training and testing set, with 100 to 5 million SNPs used to train and test the rrBLUP model. (A) all data filters applied (mapping quality, indel and repeat removal); (B) filtered mapping quality and indel removal; (C) random selection of SNPs using all data filters; (D) GP allocation accuracy calculated using data with all filters applied. The scale on the left hand vertical axis is for correlation, and the scale on the right hand vertical axis is for accuracy.

**Supplementary Figure 6.**
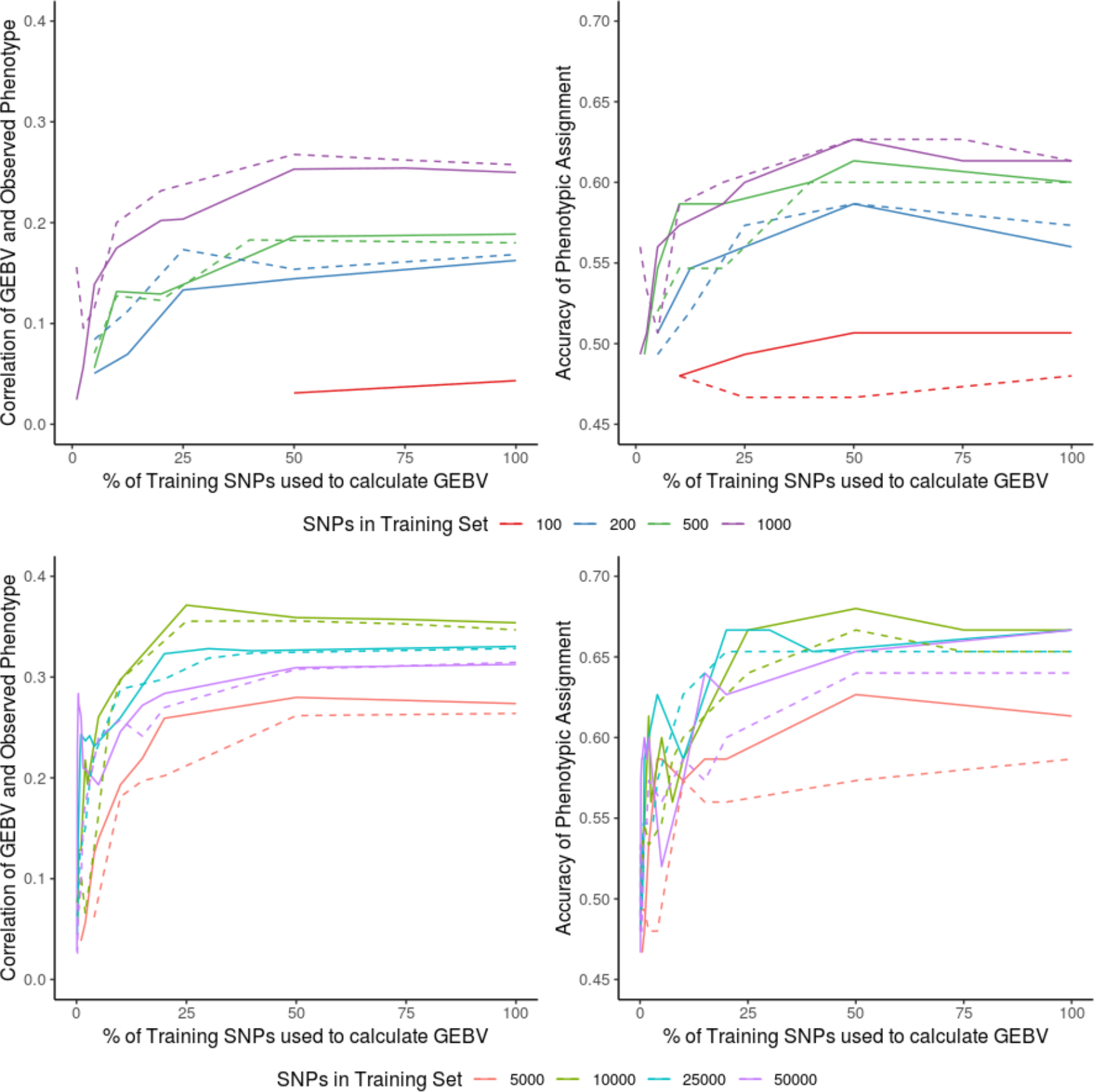
Genomic prediction using pool-seq data for training and 150 NSZ 204 individuals for testing: dashed lines show results excluding pool-seq data from provenance NSZ 204 (the test provenance) from the training dataset, whereas solid lines show results with NSZ 204 included. The left column shows correlation of observed phenotype and GEBV and the right column shows accuracy of phenotypic assignment from GEBV.

## References

1. Mitchell, R. J. et al. The potential ecological impact of ash dieback in the UK. Joint Nature Conservation Committee (2014).

2. Pautasso, M., Aas, G., Queloz, V. & Holdenrieder, O. European ash (*Fraxinus excelsior*) dieback - A conservation biology challenge. Biological Conservation (2013). doi:10.1016/j.biocon.2012.08.026

3. Sollars, E. S. A. et al. Genome sequence and genetic diversity of European ash trees. Nature (2017). doi:10.1038/nature20786

4. Gross, A., Holdenrieder, O., Pautasso, M., Queloz, V. & Sieber, T. N. *Hymenoscyphus pseudoalbidus*, the causal agent of European ash dieback. Mol. Plant Pathol. (2014). doi:10.1111/mpp.12073

5. Plumb, W. J. et al. The viability of a breeding programme for ash in the British Isles in the face of ash dieback. Plants People Planet In review

6. Mckinney, L. V. et al. The ash dieback crisis: Genetic variation in resistance can prove a long-term solution. Plant Pathology (2014). doi:10.1111/ppa.12196

7. Endler, L., Betancourt, A. J., Nolte, V. & Schlötterer, C. Reconciling differences in pool-GWAS between populations: A case study of female abdominal pigmentation in *Drosophila melanogaster*. Genetics 202, 843–855 (2016).

8. Fontanesi, L. et al. Genome-wide association study for ham weight loss at first salting in Italian Large White pigs: towards the genetic dissection of a key trait for dry-cured ham production. Anim. Genet. (2017). doi:10.1111/age.12491

9. Zhao, Y., Mette, M. F., Gowda, M., Longin, C. F. H. & Reif, J. C. Bridging the gap between marker-assisted and genomic selection of heading time and plant height in hybrid wheat. Heredity (Edinb). 112, 638–645 (2014).

10. Hayes, B. J., Visscher, P. M. & Goddard, M. E. Increased accuracy of artificial selection by using the realized relationship matrix. Genet. Res. (Camb). (2009). doi:10.1017/S0016672308009981

11. Goddard, M. E., Hayes, B. J. & Meuwissen, T. H. E. Genomic selection in livestock populations. Genet. Res. (Camb). (2010). doi:10.1017/S0016672310000613

12. Müller, B. S. F. et al. Genomic prediction in contrast to a genome-wide association study in explaining heritable variation of complex growth traits in breeding populations of Eucalyptus. BMC Genomics (2017). doi:10.1186/s12864-017-3920-2

13. Resende, J. F. R. et al. Accuracy of genomic selection methods in a standard data set of loblolly pine (*Pinus taeda* L.). Genetics (2012). doi:10.1534/genetics.111.137026

14. Schlötterer, C., Tobler, R., Kofler, R. & Nolte, V. Sequencing pools of individuals-mining genome-wide polymorphism data without big funding. Nat. Rev. Genet. 15, 749–763 (2014).

15. Stocks, J. J., Buggs, R. J. A. & Lee, S. J. A first assessment of *Fraxinus excelsior* (common ash) susceptibility to *Hymenoscyphus fraxineus* (ash dieback) throughout the British Isles. Sci. Rep. (2017). doi:10.1038/s41598-017-16706-6

16. Bakker, E. G. A Genome-Wide Survey of R Gene Polymorphisms in *Arabidopsis*. PLANT CELL ONLINE (2006). doi:10.1105/tpc.106.042614

17. Meng, Z., Ruberti, C., Gong, Z. & Brandizzi, F. CPR5 modulates salicylic acid and the unfolded protein response to manage tradeoffs between plant growth and stress responses. Plant J. (2017). doi:10.1111/tpj.13397

18. Risseeuw, E. P. et al. Protein interaction analysis of SCF ubiquitin E3 ligase subunits from Arabidopsis. Plant J. (2003). doi:10.1046/j.1365-313X.2003.01768.x

19. Baker, E. A. G. et al. Comparative Transcriptomics Among Four White Pine Species. G3 (2018). doi:10.1534/g3.118.200257

20. Kakehi, J. I. et al. Mutations in ribosomal proteins, RPL4 and RACK1, suppress the phenotype of a thermospermine-deficient mutant of arabidopsis thaliana. PLoS One (2015). doi:10.1371/journal.pone.0117309

21. Iovine, B., Iannella, M. L. & Bevilacqua, M. A. Damage-specific DNA binding protein 1 (DDB1): A protein with a wide range of functions. International Journal of Biochemistry and Cell Biology (2011). doi:10.1016/j.biocel.2011.09.001

22. Liu, Y. et al. A gene cluster encoding lectin receptor kinases confers broad-spectrum and durable insect resistance in rice. Nature Biotechnology (2015). doi:10.1038/nbt.3069

23. Hao, W., Collier, S. M., Moffett, P. & Chai, J. Structural basis for the interaction between the potato virus X resistance protein (Rx) and its cofactor ran GTPase-activating protein 2 (RanGAP2). J. Biol. Chem. (2013). doi:10.1074/jbc.M113.517417

24. Wang, S. et al. A noncanonical role for the CKI-RB-E2F cell-cycle signaling pathway in plant effector-triggered immunity. Cell Host Microbe (2014). doi:10.1016/j.chom.2014.10.005

25. Rivas-San Vicente, M. & Plasencia, J. Salicylic acid beyond defence: Its role in plant growth and development. Journal of Experimental Botany (2011). doi:10.1093/jxb/err031

26. Morita-Yamamuro, C. et al. The Arabidopsis gene CAD1 controls programmed cell death in the plant immune system and encodes a protein containing a MACPF domain. Plant Cell Physiol. (2005). doi:10.1093/pcp/pci095

27. Han, J. Y., In, J. G., Kwon, Y. S. & Choi, Y. E. Regulation of ginsenoside and phytosterol biosynthesis by RNA interferences of squalene epoxidase gene in Panax ginseng. Phytochemistry (2010). doi:10.1016/j.phytochem.2009.09.031

28. Wang, K., Senthil-Kumar, M., Ryu, C.-M., Kang, L. & Mysore, K. S. Phytosterols Play a Key Role in Plant Innate Immunity against Bacterial Pathogens by Regulating Nutrient Efflux into the Apoplast. PLANT Physiol. (2012). doi:10.1104/pp.111.189217

29. Gupta, S. K., Rai, A. K., Kanwar, S. S. & Sharma, T. R. Comparative analysis of zinc finger proteins involved in plant disease resistance. PLoS One (2012). doi:10.1371/journal.pone.0042578

30. Soll, J. & Schleiff, E. Protein import into chloroplasts. Nature Reviews Molecular Cell Biology (2004). doi:10.1038/nrm1333

31. Stief, A. et al. Arabidopsis miR156 Regulates Tolerance to Recurring Environmental Stress through SPL Transcription Factors. Plant Cell (2014). doi:10.1105/tpc.114.123851

32. Michaels, S. D. & Amasino, R. M. Memories of winter : vernalization and the competence to flower. Plant, Cell Environ. (2000). doi:10.1046/j.1365-3040.2000.00643.x

33. Liu, G., Holub, E. B., Alonso, J. M., Ecker, J. R. & Fobert, P. R. An Arabidopsis NPR1-like gene, NPR4, is required for disease resistance. Plant J. (2005). doi:10.1111/j.1365-313X.2004.02296.x

34. Gutterson, N. & Reuber, T. L. Regulation of disease resistance pathways by AP2/ERF transcription factors. Current Opinion in Plant Biology (2004). doi:10.1016/j.pbi.2004.04.007

35. Mitchell, D. A., Vasudevan, A., Linder, M. E. & Deschenes, R. J. Protein palmitoylation by a family of DHHC protein S-acyltransferases. J. Lipid Res (2006). doi: R600007-JLR200[pii]\n10.1194/jlr.R600007-JLR200

36. Li, Y., Scott, R., Doughty, J., Grant, M. & Qi, B. Protein S -Acyltransferase 14: A Specific Role for Palmitoylation in Leaf Senescence in *Arabidopsis*. Plant Physiol. (2016). doi:10.1104/pp.15.00448

37. Sharmin, S. et al. Xyloglucan endotransglycosylase/hydrolase genes from a susceptible and resistant jute species show opposite expression pattern following Macrophomina phaseolina infection. Commun. Integr. Biol. (2012). doi:10.4161/cib.21422

38. Okazawa, K. et al. Molecular cloning and cDNA sequencing of endoxyloglucan transferase, a novel class of glycosyltransferase that mediates molecular grafting between matrix polysaccharides in plant cell walls. J. Biol. Chem. (1993).

39. Sakuma, Y. et al. DNA-binding specificity of the ERF/AP2 domain of Arabidopsis DREBs, transcription factors involved in dehydration- and cold-inducible gene expression. Biochem. Biophys. Res. Commun. (2002). doi:10.1006/bbrc.2001.6299

40. Gkizi, D., Santos-Rufo, A., Rodríguez-Jurado, D., Paplomatas, E. J. & Tjamos, S. E. The β-amylase genes: Negative regulators of disease resistance for Verticillium dahliae. Plant Pathol. (2015). doi:10.1111/ppa.12360

41. Huibers, R. P., de Jong, M., Dekter, R. W. & Van den Ackerveken, G. Disease-specific expression of host genes during downy mildew infection of *Arabidopsis*. Mol. Plant. Microbe. Interact. (2009). doi:10.1094/MPMI-22-9-1104

42. Carter, C. The Vegetative Vacuole Proteome of *Arabidopsis thaliana* Reveals Predicted and Unexpected Proteins. PLANT CELL ONLINE (2004). doi:10.1105/tpc.104.027078

43. Castaño-Miquel, L. et al. SUMOylation Inhibition Mediated by Disruption of SUMO E1-E2 Interactions Confers Plant Susceptibility to Necrotrophic Fungal Pathogens. Mol. Plant (2017). doi:10.1016/j.molp.2017.01.007

44. Mur, L. A. J., Simpson, C., Kumari, A., Gupta, A. K. & Gupta, K. J. Moving nitrogen to the centre of plant defence against pathogens. Annals of Botany (2017). doi:10.1093/aob/mcw179

45. Gao, Y. et al. Two Trichome Birefringence-Like Proteins Mediate Xylan Acetylation, Which Is Essential for Leaf Blight Resistance in Rice. Plant Physiol. (2017). doi:10.1104/pp.16.01618

46. Slavov, G. T. et al. Genome-wide association studies and prediction of 17 traits related to phenology, biomass and cell wall composition in the energy grass Miscanthus sinensis. New Phytol. 201, 1227–1239 (2014).

47. Grinberg, N. F. et al. Implementation of Genomic Prediction in Lolium perenne (L.) Breeding Populations. Front. Plant Sci. 7, 1–10 (2016).

48. Spindel, J. et al. Genomic Selection and Association Mapping in Rice (Oryza sativa): Effect of Trait Genetic Architecture, Training Population Composition, Marker Number and Statistical Model on Accuracy of Rice Genomic Selection in Elite, Tropical Rice Breeding Lines. PLoS Genet. (2015). doi:10.1371/journal.pgen.1004982

49. Biazzi, E. et al. Genome-wide association mapping and genomic selection for alfalfa (Medicago sativa) forage quality traits. PLoS One 12, 1–17 (2017).

50. Bian, Y. & Holland, J. B. Enhancing genomic prediction with genome-wide association studies in multiparental maize populations. Heredity (Edinb). (2017). doi:10.1038/hdy.2017.4

51. Resende, R. T. et al. Assessing the expected response to genomic selection of individuals and families in Eucalyptus breeding with an additive-dominant model. Heredity (Edinb). (2017). doi:10.1038/hdy.2017.37

52. Hayes, B. J., Lewin, H. A. & Goddard, M. E. The future of livestock breeding: Genomic selection for efficiency, reduced emissions intensity, and adaptation. Trends in Genetics (2013). doi:10.1016/j.tig.2012.11.009

53. Pryce, J. E. & Daetwyler, H. D. Designing dairy cattle breeding schemes under genomic selection: A review of international research. Animal Production Science (2012). doi:10.1071/AN11098

54. Wientjes, Y. C. J., Veerkamp, R. F. & Calus, M. P. L. The effect of linkage disequilibrium and family relationships on the reliability of genomic prediction. Genetics (2013). doi:10.1534/genetics.112.146290

55. Clark, S. A., Hickey, J. M., Daetwyler, H. D. & van der Werf, J. H. J. The importance of information on relatives for the prediction of genomic breeding values and the implications for the makeup of reference data sets in livestock breeding schemes. Genet. Sel. Evol. (2012). doi:10.1186/1297-9686-44-4

56. Alfas, P., Lygis, V., Suchockas, V. & Bartkevičius, E. Performance of twenty four european *Fraxinus excelsior* populations in three lithuanian progeny trials with a special emphasis on resistance to *Chalara fraxinea*. Balt. For. (2011).

57. Gautier, M. et al. Estimation of population allele frequencies from next-generation sequencing data: Pool-versus individual-based genotyping. Mol. Ecol. 22, 3766–3779 (2013).

58. Bolger, A. M., Lohse, M. & Usadel, B. Trimmomatic: A flexible trimmer for Illumina sequence data. Bioinformatics (2014). doi:10.1093/bioinformatics/btu170

59. Li, H. Aligning sequence reads, clone sequences and assembly contigs with BWA-MEM. (2013).

60. Kofler, R., Pandey, R. V. & Schlötterer, C. PoPoolation2: Identifying differentiation between populations using sequencing of pooled DNA samples (Pool-Seq). Bioinformatics 27, 3435–3436 (2011).

61. Jombart, T. adegenet: a R package for the multivariate analysis of genetic markers. Bioinformatics (2008). doi:10.1093/bioinformatics/btn129

62. Wei, T. & Simko, V. Package ‘corrplot: visualization of a correlation matrix’ (v.0.84). URL https://CRAN.R-project.org/package=corrplot (2017).

63. Landis, J. R., Heyman, E. R. & Koch, G. G. Average Partial Association in Three-Way Contingency Tables: A Review and Discussion of Alternative Tests. Int. Stat. Rev. / Rev. Int. Stat. (1978). doi:10.2307/1402373

64. Storey, J. D., Bass, A. J., Dabney, A., Robinson, D. & Warnes, G. qvalue: Q-value estimation for false discovery rate control. R (2019).

65. Cingolani, P. et al. A program for annotating and predicting the effects of single nucleotide polymorphisms, SnpEff: SNPs in the genome of *Drosophila melanogaster* strain w1118; iso-2; iso-3. Fly (Austin). (2012). doi:10.4161/fly.19695

66. Waterhouse, A. et al. SWISS-MODEL: Homology modelling of protein structures and complexes. Nucleic Acids Res. (2018). doi:10.1093/nar/gky427

67. Kelley, L. A., Mezulis, S., Yates, C. M., Wass, M. N. & Sternberg, M. J. E. The Phyre2 web portal for protein modeling, prediction and analysis. Nat. Protoc. (2015). doi:10.1038/nprot.2015.053

68. Schrödinger, L. The PyMOL molecular graphics system, version 1.8. https://www.pymol.org/citing (2015).

69. Trott oleg & Arthur J. Olson. AutoDock Vina: Improving the Speed and Accuracy of Docking with a New Scoring Function, Efficient Optimization, and Multithreading. J. Comput. Chem. (2010). doi:10.1002/jcc

70. Dallakyan, S. & Olson, A. J. Small-molecule library screening by docking with PyRx. Methods Mol. Biol. (2015). doi:10.1007/978-1-4939-2269-7_19

71. Endelman, J. B. Ridge Regression and Other Kernels for Genomic Selection with R Package rrBLUP. Plant Genome J. (2011). doi:10.3835/plantgenome2011.08.0024

72. Endelman, J. B. & Jannink, J.-L. Shrinkage Estimation of the Realized Relationship Matrix. Genes|Genomes|Genetics (2012). doi:10.1534/g3.112.004259

